# IFNγ-Expressing Myeloid Cells Localize within Lipoproteinosis during Drug-Associated Pulmonary Alveolar Proteinosis occurring in Systemic Juvenile Idiopathic Arthritis

**DOI:** 10.1101/2024.11.22.624756

**Authors:** Alea Delmastro, Candace C. Liu, Xiao-Wen Ding, Serena Y. Tan, Inna Averbukh, Marc Bosse, Timothy J. Keyes, Surbhi Sharma, Gail Deutsch, Michael Angelo, Vivian E. Saper, Elizabeth D. Mellins, Erin F. McCaffrey

## Abstract

In the United States, approximately one in 1000 children are diagnosed with the autoinflammatory disease, Juvenile Idiopathic Arthritis (JIA). A subset of JIA cases manifests as Systemic JIA (sJIA), which is characterized by joint pain, fevers, rashes, and systemic inflammation. Severe pulmonary complications have not historically been associated with sJIA. Since 2010, inhibitors of interleukin-1 and interleukin 6 (IL-1i/IL-6i) are the recommended course of treatment for sJIA, yet recently studies show evidence of a severe drug hypersensitivity reaction implicating these medications in a subset of those treated. With this reaction, sJIA patients can develop severe lung disease, including pulmonary alveolar proteinosis (PAP). As this drug-associated lung disease has only recently been identified, the etiology of sJIA drug-associated PAP (sJIA-daPAP) is poorly understood. We used multiplexed ion beam imaging by time-of-flight (MIBI-TOF) to define the cellular immune infiltrate and describe pathological features of PAP in sJIA-daPAP patients. We found an enrichment of eosinophils, neutrophils, and M2 macrophages within regions of lipoproteinosis. These enriched subsets all upregulate IFNγ within lipoproteinosis, a signature specific to sJIA-daPAP samples compared to non-sJIA-PAP samples. In a cellular neighborhood analysis, we identified that eosinophils, neutrophils and M2 macrophages frequently co-localize within the same cellular microenvironment, especially in lipoproteinosis regions. Therefore, this spatial coordination may be involved in clearance or persistence of lipoproteinosis in sJIA-daPAP. This study provides a comprehensive overview of sJIA-daPAP immune pathology and suggests cellular mechanisms that drive inflammation in sJIA patients experiencing pulmonary complications associated with delayed drug hypersensitivity during IL-1i/IL-6i treatment.

## Introduction

Systemic juvenile idiopathic arthritis (sJIA) is an autoinflammatory illness that predominantly affects the pediatric population, with adult-onset Still disease as its adult counterpart. It is characterized by quotidian fever, arthritis or arthralgias, evanescent rash, hepatosplenomegaly, lymphadenopathy, serositis (including pleuritis), and elevated inflammatory markers^1^. Severe systemic inflammation includes the risk of potentially fatal macrophage activation syndrome (MAS)^2,3^. Since 2010, treatment with inhibitors of interleukin-1 and interleukin 6 (IL-1i and IL-6i, respectively) has become common and has significantly improved Still disease prognosis^3,4^.

Historically, pulmonary involvement has not been a feature of Still disease^4^. During IL-1i/IL-6i treatment, however, some patients develop an unusual, severe pulmonary disease, including diffuse lung disease with or without pulmonary hypertension^3,4^. In patients with this lung disease, high levels of IL-18 were observed in serum and in bronchoalveolar lavage (BAL), in which neutrophils were predominant^5^. Lung biopsy samples from these patients revealed a patchy but extensive form of pulmonary alveolar proteinosis (PAP)^5^. Generally, in PAP, alveoli become filled with granular, eosinophilic material, including proteinaceous material derived from pulmonary surfactant, phospholipids, and cholesterol^6^. PAP can coincide with endogenous lipoid pneumonia (ELP), when fat-filled vacuolated alveolar macrophages (“foamy macrophages”) fill the alveoli^3,5^.

PAP is a rare pulmonary syndrome characterized by the presence of alveolar proteinosis on histology^7^. The specific underlying causes vary in different types of PAP, with congenital, autoimmune, and secondary etiologies, which affect either surfactant metabolism in type II pneumocytes or macrophage clearance^8^. The specific underlying causes vary in different types of PAP^8^. Congenital PAP is caused by mutations in genes, such as *SFTPB, SFTPC,* and *ABCA3*, which play roles in surfactant metabolism. Primary, autoimmune PAP is the most common form, occurs in adults, and accounts for 90% of PAP cases overall. It involves the disruption of granulocyte-macrophage colony-stimulating factor (GM-CSF) pathway by anti-GM-CSF autoantibodies^8^. A much rarer congenital form of primary PAP disrupts the GM-CSF signaling pathway through mutation of *CSF2RA or CSF2RB,* parts of the GM-CSF receptor^8^. Secondary PAP is associated with a variety of conditions, such as immunodeficiency or hematologic disorders as well as chronic infections, that reduce the number or function of alveolar macrophages^8^. Recent evidence suggests secondary PAP can be drug-induced^8,9^. Drug-associated PAP cases also appear in sJIA patients during treatment with IL-1i and/or IL-6i^3,10^; these particular PAP cases will now be referred as sJIA drug-associated PAP (sJIA-daPAP).

Importantly, sJIA-daPAP appears to be novel. Patients lack genetic or autoantibody-mediated dysfunction of the GM-CSF pathway, and BAL nucleated cell counts show reduced levels of lipid-laden macrophages^5^. PAP samples from sJIA patients frequently show persistent lung injury, as evidenced by lobular remodeling with fibrosis, pulmonary vascular involvement (arterial wall thickening), lymphoplasmacytic infiltration (CD4+ T predominant), and cholesterol clefts^5,11^. Cholesterol clefts reflect cholesterol esters that become organized after release from injured cells^12^. In a recent study, we found that, at the ultrastructural level, the acellular material and alveolar macrophages contained multicellular structures. The pulmonary findings were preceded by features that met criteria for DReSS (drug reaction with eosinophilia and systemic symptoms) by RegiSCAR scoring^11^. Subsequently, we found that 80% of those satisfying RegiSCAR criteria carried one of a subset of HLA-DRB1*15 alleles (i.e.,15:01,15:03,15:06) that share similar peptide binding specificity and are tightly linked to DRB5*01:01^11^. In addition, the serum proteome profile from sJIA-daPAP patients differs from that of primary PAP patients, with sJIA-daPAP serum enriched for sICAM5, MMP7, and CCL17, compared to serum from primary PAP^13–16^. These markers also are associated with other inflammatory or fibrotic forms of pulmonary disease, including idiopathic pulmonary fibrosis and DReSS^17,18^. Taken altogether, these observations suggest that sJIA-daPAP exhibits distinct pathology compared to other forms of PAP.

Currently, the etiology of sJIA-daPAP is poorly understood. The objective of this study was to comprehensively assess the immune pathology of sJIA-daPAP by investigating the immunophenotype, functional profile, cellular interactions, and spatial distribution of immune cells in sJIA-daPAP compared to uninvolved healthy lung tissue from the same biopsy. We employed multiplexed ion beam imaging by time-of-flight (MIBI-TOF) to spatially map the cellular infiltrate and pathological features of PAP in archival pulmonary tissue from a cohort of pediatric patients with sJIA-daPAP^19,20^. In doing so, we revealed both enrichment and unique functional profiles of several myeloid cell populations, including IFNγ-expressing eosinophils, within lipoproteinosis regions and identified features of sJIA-daPAP. We also employed spatial analyses to define the cellular microenvironments (MEs) that may be involved in clearance or persistence of lipoproteinosis. This study provides a comprehensive census of immune pathology during sJIA-daPAP and suggests mechanisms in this emerging complication affecting a subset of sJIA patients treated with IL-1i/IL-6i treatment.

## Results

### Study Design, Technological Overview, and Analytical Outline

To study the immunological features of the PAP lung disease arising in sJIA patients, we assembled a cohort of pulmonary tissue samples from pediatric patients from six institutions^3,5^. In total, we procured 14 pulmonary biopsy specimens, comprising sJIA-daPAP (n = 12 specimens from 9 patients; **Table S1**) and non-sJIA-PAP (n = 2 specimens from 2 patients; **Table S2**). sJIA-daPAP lung tissue showed PAP disease in patients diagnosed with sJIA and undergoing IL-1i and/or IL-6i treatment at the time of biopsy. Each sJIA patient, retrospectively, had scored as having DReSS by RegiSCAR, implicating these medications. Non-sJIA-PAP showed PAP disease in patients without sJIA: one with GATA2-deficiency/PAP^13,21^ and one with COPA syndrome/PAP^22^. A team of pathologists selected several fields-of-view (FOVs) from each biopsy specimen to account for interpatient heterogeneity and to sample regions of uninvolved lung adjacent to the PAP pathology (**Figures 1a-b**).

**Figure 1:**
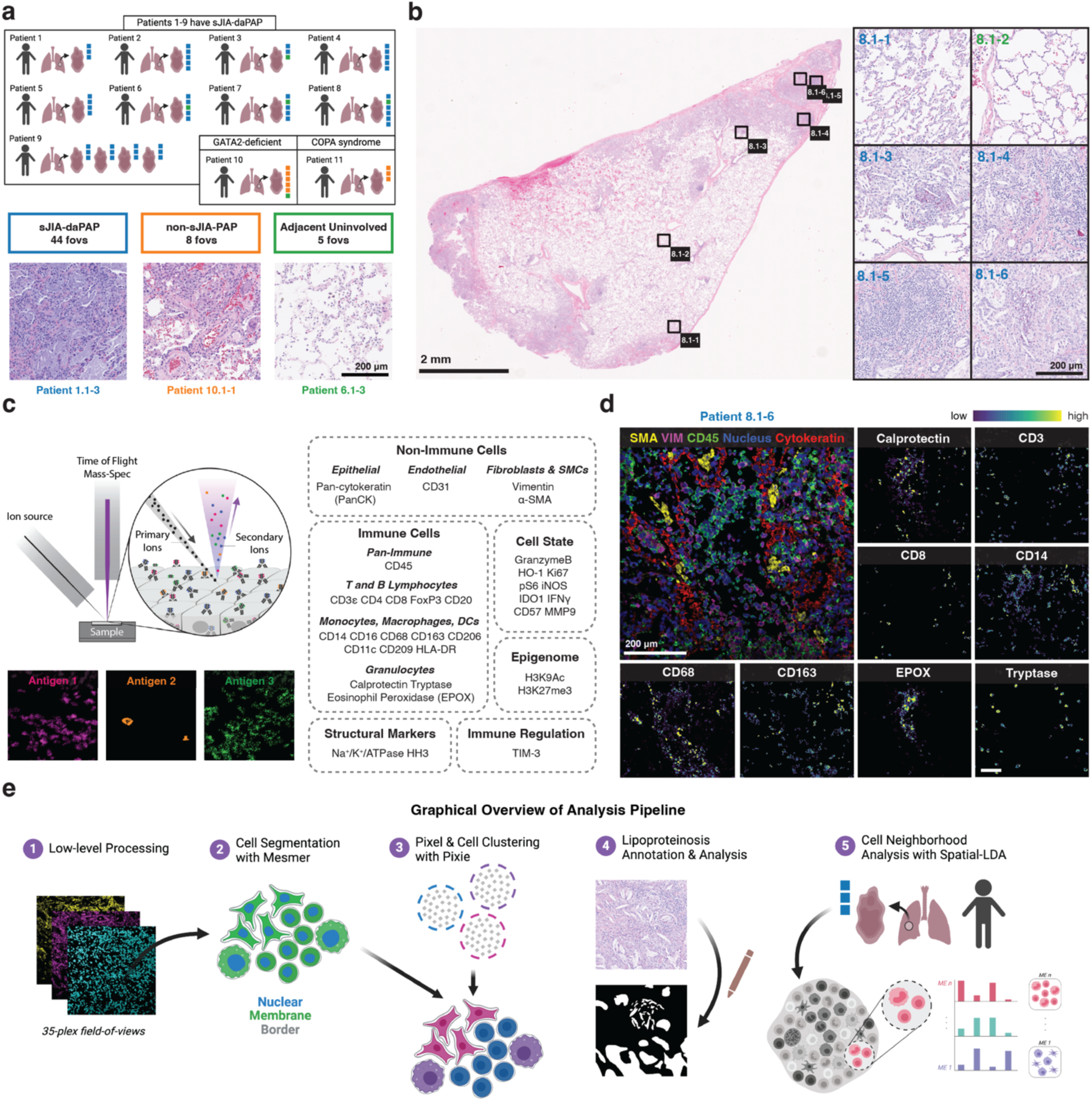
Study design and analytical overview. **(a)** Cohort characteristics, including the number of specimens per patients and the number of 500 µm x 500 µm field-of-views (FOVs) acquired per specimen, color-coded by disease category. Below includes representative hematoxylin and eosin (H&E) stained FOVs for each of the three disease categories and lists the total number of FOVs acquired per category. Per patient clinical characteristics are provided in Tables S1 and S2. **(b)** On the left, one representative H&E-stained specimen along with the locations of the FOVs acquired. On the right, the FOVs from the specimen on the left, color-coded by their disease category assignment. **(c)** Graphical illustration of MIBI-TOF methodology (right) and the list of markers included in the imaging panel. **(d)** One representative MIBI-TOF overlay, demonstrating major lineage markers: SMA (yellow), vimentin (magenta), CD45 (green), HH3 (blue), and PanCK (red). Smaller images denote expression patterns for the listed phenotypic markers from the same FOV as overlay. **(e)** Graphical overview of the entire analytical pipeline applied to the data set as part of this study.

We used MIBI-TOF to image the selected FOVs after staining with a 36-plex panel of metal-labeled antibodies (**Figures 1c-d and S1, Table S3**)^20,23^. This panel included markers to identify major immune and non-immune cell lineages, including lymphocytes, macrophages, granulocytes, stroma, and epithelium. We also included functional markers to delineate cellular activation, immune regulation, and epigenetic state. After cell identification and classification, we identified lipoproteinosis and cellular neighborhoods with our custom analytical pipeline (**Figure 1e**).

### Defining the cellular composition of sJIA-daPAP with respect to adjacent uninvolved lung and non-sJIA-PAP

We first sought to define the cellular phenotypes uniquely associated with sJIA-daPAP and non-sJIA-PAP. We identified 21 unique cell subsets, including several monocyte-lineage subsets (**Figure 2a**)^24,25^. M2 macrophages, which expressed CD68, CD163, and CD206 but lacked CD209, encompassed alveolar, infiltrating, and foamy macrophages, as confirmed by histological review^26^. We then mapped all 21 cellular phenotypes back to their spatial position in each FOV to generate cell phenotype maps (CPMs) for downstream compositional and spatial analyses (**Figures 2b and S2)**. While lung epithelium and mesenchymal cells unsurprisingly made up a large percentage of the cells present throughout all the FOVs, a significant immune infiltrate was evident, including CD4+ T cells and neutrophils (**Figure 2a**). We also annotated 13 giant cells, which had a phenotypic profile similar to macrophages and appeared foamy histologically (**Figure 2a**).

**Figure 2:**
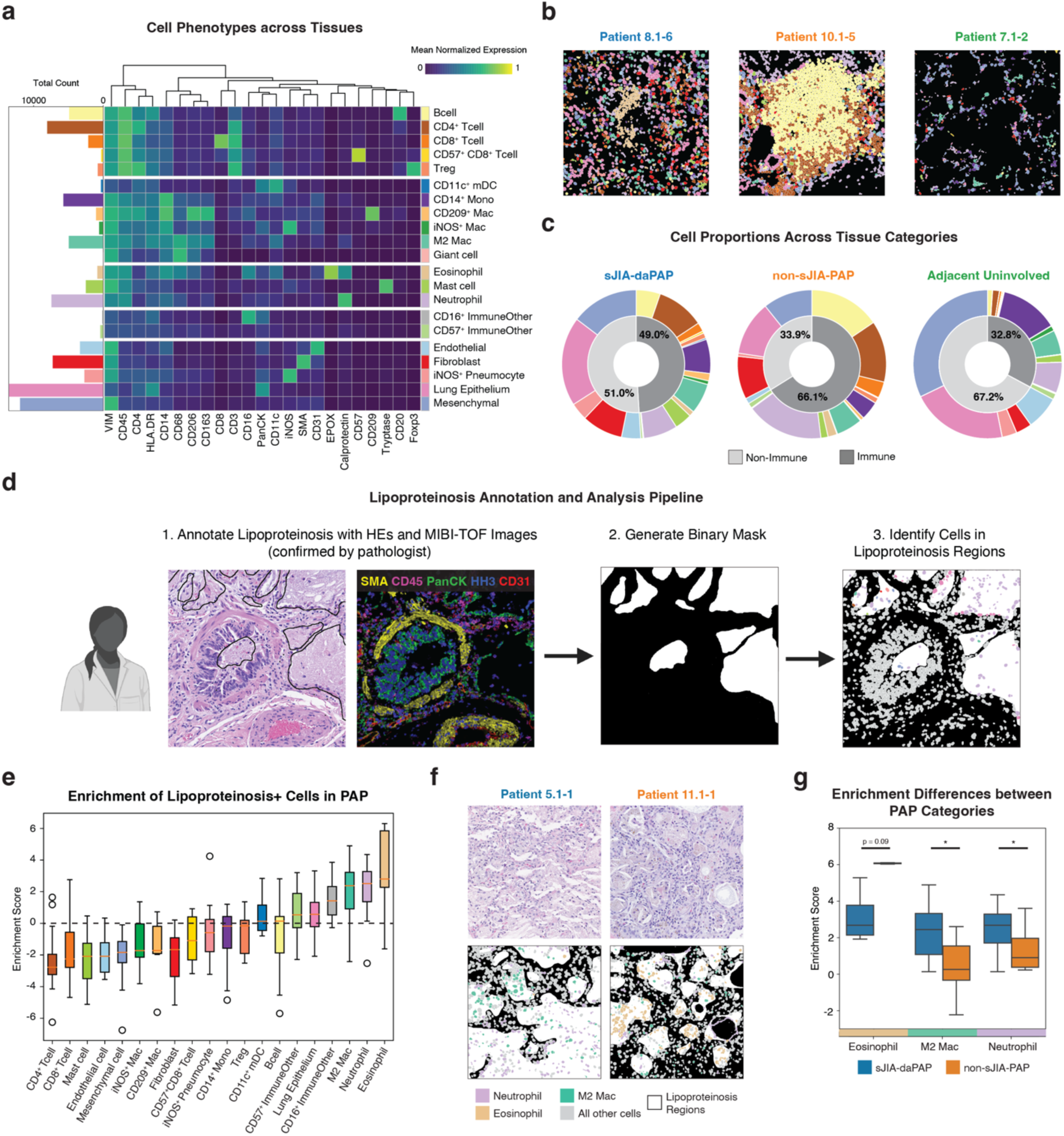
Cellular landscape in PAP samples and within lipoproteinosis. **(a)** Cell lineage assignments based on normalized expression of lineage markers (heatmap columns). Bar plot (left) denotes absolute abundance. Rows are ordered by phenotype group (lymphocytes, myeloid cells, granulocytes, immune “other”, and non-immune), whereas columns are hierarchically clustered (Euclidean distance, average linkage). **(b)** Representative cell phenotype maps (CPMs). **(c)** Cell subset (outer donut) and immune vs. non-immune (inner donut) proportions by disease group. Per patient cell subset proportions are provided in Tables S1 and S2. **(d)** Conceptual overview of lipoproteinosis annotation and analysis. **(e)** Enrichment score distribution of lipoproteinosis+ cells by subset, ordered by median. The enrichment score is defined as the log2 ratio of the sum of lipoproteinosis+ cells over the sum of lipoproteinosis-cells for each cell subset. **(f)** Representative H&Es (top row) and CPMs (bottom row), colored only for top three enriched subsets in lipoproteinosis, overlaid with lipoproteinosis mask. **(g)** Enrichment score comparisons between sJIA-daPAP and non-sJIA-PAP FOVs. All P values were calculated with a Student’s t-test (two tailed) (*P < 0.05).

We next evaluated how cellular composition varied with clinical status and disease involvement. We broke down the cell proportions by their study category – sJIA-daPAP, non-sJIA-PAP, and adjacent uninvolved (**Figures 2c and S3a**) – as well as by FOV (**Figure S3b**) and by patient (**Tables S1 and S2**). sJIA-daPAP and non-sJIA-PAP FOVs contained higher proportions of major immune populations, such as CD4+ T cells and B cells, compared to adjacent uninvolved lung tissue. The giant cells we identified all originated from non-sJIA-PAP samples: one across the GATA2-deficient FOVs and twelve across the COPA syndrome FOVs, which may indicate functional differences in lipoproteinosis clearance compared to sJIA-daPAP. The low sample size of the non-sJIA-PAP group precluded deep comparison of patient-level features. However, to detect statistically significant differences in cell abundances between involved and adjacent uninvolved regions within sJIA-daPAP biopsy specimens, we applied binomial mixed modeling. This analysis revealed an abundance of multiple immune cell subsets, including CD4+ T cells, B cells, neutrophils, eosinophils and M2 macrophages, in sJIA-daPAP involved regions compared to adjacent uninvolved regions from the same specimen (**Figure S4**). As expected, sJIA-daPAP involved regions had a decrease in the relative abundance of lung epithelial cells, which are highly prevalent in healthy lungs. Our results are consistent with a substantial and diverse immune infiltrate localized to the site of PAP pathology (**Figure 2c**).

### Characterizing the cellular landscape within lipoproteinosis regions

The defining histological feature of PAP is surfactant accumulation or lipoproteinosis. Considering this, it is important to know which cell types are specifically recruited to these sites. To achieve this, we spatially mapped all cells in sJIA-daPAP and non-sJIA-PAP diseased regions and distinguished between cells within lipoproteinosis regions (LPRs) and those surrounding these regions without lipoproteinosis (non-LPRs) (**Figures 2d and S2**). The three most enriched cell subsets in LPRs were eosinophils, neutrophils, and M2 macrophages (**Figure 2e**). M2 macrophages are known to produce the chemokine thymocyte and activation-regulated chemokine (TARC)/CCL17 and recruit eosinophils^27^. We visually confirmed these findings in representative CPMs overlaid with their respective lipoproteinosis mask (**Figure 2f)**. In addition to lung epithelium, which line the alveoli and act as a physical barrier, these three immune subsets made up a large proportion (∼46%) of the cells within LPRs (**Figure S5a**). Interestingly, M2 macrophages and neutrophils were more enriched in sJIA-daPAP than non-sJIA-PAP lipoproteinosis regions (**Figure 2g**). Taken together, we found that lipoproteinosis contained a niche cellular landscape, primarily consisting of infiltrating innate immune cells.

### Eosinophils in lipoproteinosis regions are associated with an IFNγ signature in sJIA-daPAP samples

We were next interested in describing the functional profile of immune cells in these LPRs. We first performed a differential expression analysis and found that cells in LPRs upregulated the proinflammatory cytokine IFNγ, HLA-DR, Ki67, and calprotectin and downregulated iNOS and CD57 compared to their non-LPR counterparts (**Figure 3a**). When we compared differential expression of these functional markers in LPRs between the sJIA-daPAP and non-sJIA-PAP, we found that IFNγ was exclusively upregulated in sJIA-daPAP, and not in non-sJIA-PAP LPRs (**Figure S5b**). Among the cell populations enriched in LPRs, we found that eosinophils, neutrophils, and M2 macrophages had higher IFNγ expression in sJIA-daPAP samples than in non-sJIA-PAP samples (**Figure 3b**). This relationship was not present in the next most enriched subsets: neither lung epithelium nor CD16+ immune other cells (**Figures S5c-d**), suggesting that the IFNγ signature in the sJIA-daPAP specimens is driven by eosinophils, neutrophils, and M2 macrophages. In sJIA-daPAP samples, the mean IFNγ intensity of eosinophils in LPRs was higher than that of both neutrophils (Student’s t-test, p < 0.0001) and M2 macrophages (Student’s t-test, p < 0.0001) in LPRs (**Figure 3b**). Visually, eosinophils expressed significant levels of IFNγ, especially in sJIA-daPAP specimens (**Figure 3c**). These IFNγ-expressing eosinophils also co-expressed TIM-3, a checkpoint molecule, and HLA-DR, the receptor necessary for antigen-presentation (**Figure 3d**)^28^. This functional signature was more elevated within the sJIA-daPAP-involved regions than the non-sJIA-PAP-involved regions (**Figures 3d-e**). Our data suggest that eosinophils are activated and proinflammatory in sJIA-daPAP samples given their high expression of IFNγ, TIM-3, and HLA-DR^29,30^.

**Figure 3:**
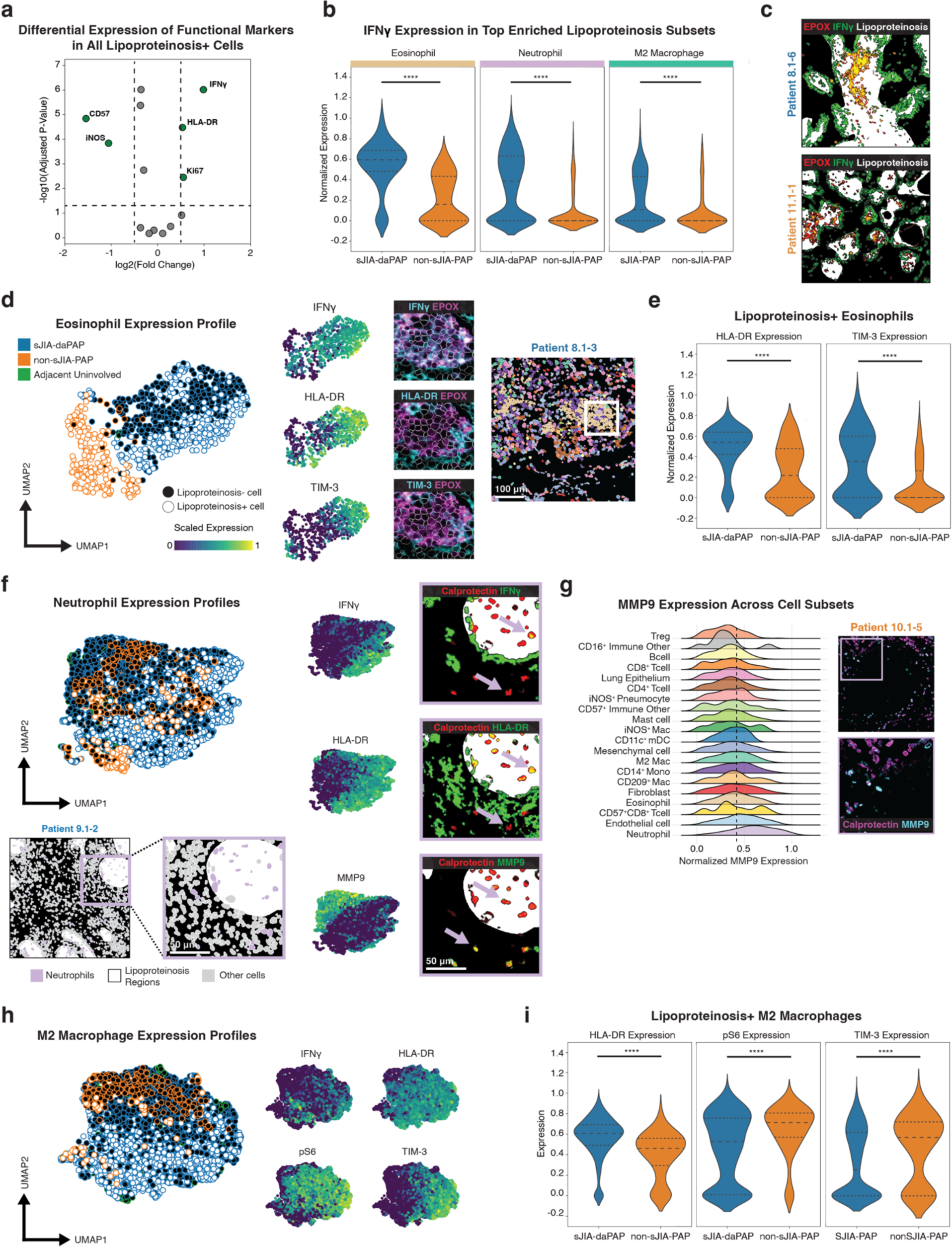
Functional profile of cells enriched in lipoproteinosis regions. **(a)** Differential expression of lipoproteinosis+ cells. Significance threshold of absolute value of log2FC > 0.5 and adjusted p-value of 0.05, with Benjamini/Hochberg multiple hypothesis correction applied. **(b)** Comparison of IFN𝛾 expression by eosinophils, neutrophils, and M2 macrophages within lipoproteinosis regions between sJIA-daPAP and non-sJIA-PAP samples. **(c)** Representative overlays of EPOX (red), IFN𝛾 (green), and lipoproteinosis (white). Yellow indicates co-expression of EPOX and IFN𝛾. **(d)** UMAP of eosinophils (left). Each dot represents one cell, outlined with the study category color and filled with white (lipoproteinosis+) or black (lipoproteinosis-). Feature plots for IFN𝛾, HLA-DR, and TIM-3 are highlighted, and representative expression overlays are provided adjacently: EPOX (magenta) and functional marker (cyan). The cell phenotype map (CPM; right) represents the FOV from which the expression overlays were obtained. **(e)** Comparison of HLA-DR and TIM-3 expression by eosinophils within lipoproteinosis regions between sJIA-daPAP and non-sJIA-PAP samples. **(f)** UMAP of neutrophils (top left). CPM from a representative FOV, overlaid with lipoproteinosis mask and colored for only neutrophils, denotes lipoproteinosis region inset (bottom left). Feature plots for IFN𝛾, HLA-DR, and MMP9 are highlighted, and representative expression overlays are provided adjacently: Calprotectin (red), functional marker (green), and lipoproteinosis (white). Yellow indicates co-expression of Calprotectin and functional marker. **(g)** Histogram of MMP9 expression (expression > 0) for each cell subset in involved and adjacent uninvolved PAP regions (left) with representative expression overlay and inset of Calprotectin (magenta) and MMP9 (cyan) (right). **(h)** UMAP of M2 macrophages (left) and feature plots for IFN𝛾, HLA-DR, pS6, and TIM-3 (right). **(i)** Comparison of HLA-DR, pS6, and TIM-3 expression by M2 macrophages within lipoproteinosis regions between sJIA-daPAP and non-sJIA-PAP samples. All P values were calculated with a Student’s t-test (two tailed) (****P < 0.0001).

### Neutrophils and M2 macrophages have reduced capacity to clear lipoproteinosis within pathology

In addition to increased IFNγ expression in eosinophils, we found that neutrophils in LPRs upregulated IFNγ as well (**Figures 3b, 3f, and S5d**). Given that IFNγ can also induce HLA-DR expression on neutrophils, it is unsurprising that we found neutrophils in LPRs highly expressed HLA-DR (**Figure 3f**)^31^. However, IFNγ+ neutrophils lacked expression of MMP9, an enzyme heavily involved in matrix degradation^32^. Neutrophils are the primary source of MMP9 in these specimens (**Figure 3g**); yet they predominately expressed MMP9 outside of lipoproteinosis, in both sJIA-daPAP and non-sJIA-PAP samples (**Figures 3f and S5d-e**). This suggests that, while neutrophils may be recruited to the site of pathology for the purpose of wound healing and lipoproteinosis clearance, they shift to a more inflammatory state once within a LPR.

We saw a similar shift away from lung repair in M2 macrophages. Like eosinophils and neutrophils, M2 macrophages within LPRs upregulated IFNγ and HLA-DR (**Figures 3h and S5d**). They also expressed high levels of pS6 and TIM-3. Interestingly, M2 macrophages in LPRs expressed lower levels of pS6 and TIM-3 in sJIA-daPAP compared to those in non-sJIA-PAP (**Figure 3i**). On macrophages, TIM-3 can participate in inflammatory cell activation. The co-expression of pS6, HLA-DR, IFNγ, and TIM-3 suggests that M2 macrophages engage in antigen presentation, possibly driving inflammatory signaling pathways and regulating immune responses in this pathologic environment^33^.

### Immune cells organize into microenvironments with distinct biological programs and spatial distributions in PAP

Cell-to-cell interactions dictate immune responses and can provide information on how pathological features, such as lipoproteinosis, reinforce or restrict cellular organization. To decouple the relationship between cellular interactions, functional state, presence of lipoproteinosis, and spatial microenvironment, we applied cellular neighborhood analysis. Briefly, we employed spatial-Latent Dirichlet Allocation (spatial-LDA) to identify recurrent patterns of spatial association between cell types^34^. Altogether we found nine distinct MEs across all FOVs (**Figures 4a-b, S2, and S6a**).

**Figure 4:**
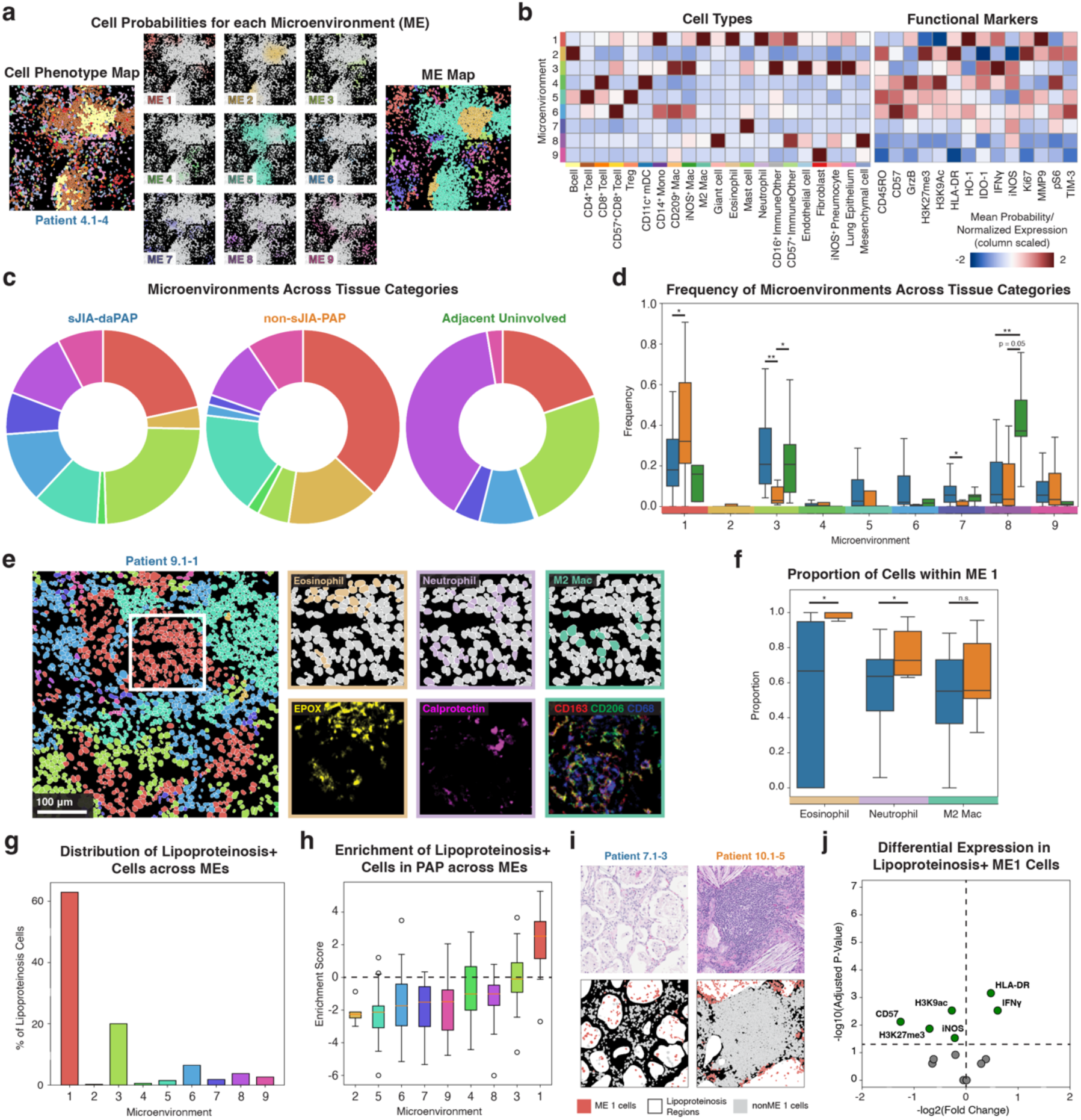
Spatial-LDA reveals lipoproteinosis niche. **(a)** Cell probability map (left), max probability microenvironment (ME) map (right), and ME probability for nine MEs (middle, scaled 0 to 1) for a representative FOV. **(b)** Heatmap of ME preferences for all subsets (standardized mean ME loading) and mean normalized expression of functional markers by assigned cells. Columns are ordered by phenotype group (lymphocytes, myeloid cells, granulocytes, immune “other”, and non-immune) and alphabetically for functional markers, whereas columns ordered numerically. **(c)** ME proportions by study group. Per patient microenvironment proportions are provided in Tables S1 and S2. **(d)** Frequency distribution comparison of each ME across the three study categories. **(e)** Representative ME map (left) with inset CPMs (top right), colored for only eosinophils, neutrophils, and M2 macrophages, respectively, and associated expression images (bottom right) for their phenotypic markers. **(f)** Frequency distribution of eosinophils, neutrophils, and M2 macrophages in ME1 out of all MEs between sJIA-daPAP and non-sJIA-PAP FOVs. **(g)** Proportion of cells within lipoproteinosis, broken down by their associated ME. **(h)** Enrichment score distribution of lipoproteinosis+ cells by ME, ordered by median. The enrichment score is defined as the log2 ratio of the sum of lipoproteinosis+ cells over the sum of lipoproteinosis-cells for each microenvironment. **(i)** Representative H&Es (top row) and ME maps (bottom row), colored only for ME1 cells, overlaid with lipoproteinosis mask. **(j)** Differential expression of lipoproteinosis+ ME1 cells. Significance threshold of absolute value of log2FC > 0 and adjusted p-value of 0.05, with Benjamini/Hochberg multiple hypothesis correction applied. All P values were calculated with a Student’s t-test (two tailed) (*P < 0.05; **P < 0.01).

To understand the context of each of these MEs, we evaluated their cellular content, functional marker expression, and distribution across all tissue groups and by patient (**Figures 4b-d, Tables S1 and S2**). ME 1 was more frequently present in non-sJIA-PAP regions compared to both sJIA-daPAP regions and adjacent uninvolved regions (**Figures 4c-d**). Notably, ME 1 was highly enriched with many previously highlighted cell subsets: eosinophils, neutrophils, and M2 macrophages (**Figure 4b**). Upon visual inspection, we saw that eosinophils, neutrophils, and M2 macrophages make up a large proportion of the cells within ME 1 (**Figures 4e and S6b**). The three subsets were all mostly associated with ME 1, yet there was some variability with affiliation; the average proportion of neutrophils and eosinophils that belong to ME 1 in sJIA-daPAP regions was less than that in non-sJIA-PAP regions (**Figure 4f**). Within ME 1, we saw an upregulation of IFNγ, IDO-1, TIM3, and MMP9 (**Figure 4b**). As described previously, MMP9 was predominantly expressed by neutrophils (**Figure 3g**). The enrichment of eosinophils and the upregulation of IFNγ and TIM-3 in ME 1 complemented our prior analysis of eosinophil states (**Figures 3d and 4b**).

Conversely, ME 3 was more prevalent in sJIA-daPAP regions compared to non-sJIA-PAP regions (**Figure 4d**). ME 3 primarily consisted of CD209+ macrophages, iNOS+ macrophages, endothelium, lung epithelium, and pneumocytes (**Figure 4b**). This ME had many “normal lung” phenotypes, such as lung epithelium and endothelial cells. Supporting this, ME 3 also made up a large frequency of the MEs present in adjacent uninvolved FOVs (**Figures 4c-d**). Along with this, ME 8 was most prevalent in adjacent uninvolved lung and represented a “normal lung” phenotype, given its enrichment of mesenchymal cells and lack of a pro-inflammatory signature.

We also observed that ME 7, an environment enriched for mast cells and upregulation of functional markers IDO-1, iNOS, and pS6, was significantly more abundant in sJIA-daPAP compared to non-sJIA-PAP regions (**Figures 4b and 4d**). This suggests that, while sJIA and non-sJIA share many similar cellular attributes, the cell populations in these samples organize in different ways and can adopt unique functional states.

To understand how lipoprotein may influence cellular MEs, we analyzed how the MEs are localized within LPRs. ME 1 comprised over 60% of the lipoproteinosis+ cells in both sJIA-daPAP and non-sJIA-PAP, compared to less than 25% of all cells (**Figures 4g and S6c**). To further support this, we quantified the enrichment of ME cells within regions of lipoproteinosis relative to the surrounding area and found that, indeed, ME 1 cells were enriched in regions of lipoproteinosis (**Figure 4h**). Out of all MEs, ME 1 was the only ME positively enriched in lipoproteinosis regions. The representative FOVs in **Figure 4i** validate that ME 1 was the most prominent ME within these regions of lipoproteinosis, for both sJIA-daPAP and non-sJIA-PAP samples. In juxtaposition to **Figure 4f**, eosinophils, neutrophils, and M2 macrophages in LPRs were all almost entirely affiliated with ME 1, for both sJIA-daPAP and non-sJIA-PAP samples (**Figure S6d**). Performing differential expression on ME 1 cells, we found that ME 1 cells in LPRs upregulated HLA-DR and IFNγ, confirming the signature we observed among eosinophils, neutrophils, and M2 macrophages in LPRs (**Figure 4j**). We also observed downregulation of CD57 and the epigenetic markers, H3K9ac and H3K27me3, suggesting that lipoproteinosis can induce an epigenetic change in immune cells; however, more samples are necessary to investigate this impact further.

Sites of chronic inflammation, for example in the lung, can lead to the development of ectopic lymphoid organs, also known as tertiary lymphoid structures (TLS)^35^. We acquired and analyzed five TLSs from three patients: patients 4 and 8 (sJIA-daPAP) and patient 10 (non-sJIA-PAP, GATA2 mutation). We investigated which lymphocytic cell populations and MEs were involved in these TLSs and how they were organized within PAP regions. **Figure S7a** depicts one such TLS and the individual lymphocyte cells present within this representative FOV. We found that TLSs primarily consisted of populations from MEs 2, 4, and 5. Among all PAP involved regions, CD4+ T cells and B cells were present at the highest frequencies (**Figure S7b**). Even though CD8+ T cells were more prevalent compared to Tregs (**Figure 2a**), there was no statistical significance between the frequency of Tregs and CD8+ T cells within TLS. Like germinal centers found in lymph nodes, B cells within these TLSs tend to be heavily compartmentalized. These B cells also made up the majority of cells within ME 2 (**Figure 4b**). Similarly, CD4+ T cells tended to primarily be concentrated within a single ME, specifically ME 5; however, there was relatively homogenous mixing of other T cell subsets within the ME 5 and the TLS, including CD8+ T cells, Tregs, and scarcely CD57+CD8+ T cells. Lastly, we considered ME 4 to be part of the TLS because of its enrichment in CD8+ T cells and its proximity to other lymphocytic MEs (**Figures 4b and S7a**). When examining the functional profile of the cell subsets within TLSs, we found that Tregs were the most proliferative and, unsurprisingly, immunosuppressive based on increased Ki67 and IDO-1 expression, respectively (**Figure S7c**). Since only five TLSs were acquired in this dataset, more samples are necessary to make statistical analyses. Future studies on the role of TLSs in PAP pathogenesis would be instrumental in understanding whether these structures are beneficial in clearing lipoprotein and reducing inflammation.

## Discussion

This study provides a novel and comprehensive analysis of the cellular and spatial landscape in sJIA-daPAP lung tissue. By employing MIBI-TOF imaging with a 36-plex panel of metal-labeled antibodies, we defined the phenotypes and functional profiles of immune cells within sJIA-daPAP histopathology. Moreover, we identified a unique IFNγ signature of sJIA-daPAP driven by myeloid cells co-localized within regions of lipoproteinosis. This analysis is the first such study of a PAP pathology and, here, occurring in the novel DReSS-associated setting^10^. In each of the nine sJIA-daPAP patients, we found that cellular features are similar regardless of IL-1i/IL-6i drug choices, HLA-DRB1*15, duration of IL-1i/IL-6i exposure, or additional immune suppressive treatment at the time of lung biopsy. Our findings may help uncover the mechanism of drug-associated PAP in sJIA patients and pave the way to clinical utility by comparing PAP signatures in other settings.

Our analysis revealed an infiltration of CD4^+^ T cells and B cells across involved regions in both sJIA-daPAP and non-sJIA-PAP. We demonstrate an overall abundance in CD4+ T cells, CD8+ T cells, and B cells in sJIA-daPAP involved regions compared to adjacent uninvolved regions. The presence of these lymphocyte subsets, particularly the predominance of CD4^+^ T cells, corroborates and expands our knowledge of immune cell infiltration in sJIA-daPAP^5^. In a study involving GM-CSF deficient mice, a model for autoimmune PAP, these mice exhibited a domination of B lymphocytes in pulmonary peribronchovascular regions^36^. These findings collectively suggest lymphocytic infiltration is a common feature of PAP. The pattern of lymphocytic infiltration may vary across different types of PAPs^37^, depending on the anatomical location examined. In non-drug-associated PAPs, lymphocytic infiltration has been observed in pulmonary alveoli, terminal bronchioles, and the walls of bronchioles but does not extend into the interalveolar septa^38,39^. Of note, lymphocytes not only infiltrated these areas of PAP pathology but, in three patients here, we observed that they organized into tertiary lymphoid structures. The role of these structures during PAP is not well characterized. While we did not delineate the differences in lymphocyte abundance across histological locations in sJIA-daPAP and non-sJIA-PAP involved regions, future studies should focus on comparing PAP subtypes based on their anatomical locations within the lung and investigate how tertiary lymphoid structures contribute to the modulation of inflammation.

The present study highlights the accumulation and co-localization of M2 macrophages, along with neutrophils and eosinophils, in the lipoproteinosis microenvironment of PAP. Macrophages are crucial mediators in the pathogenesis of PAP and may exhibit impaired prevalence, function, or both^6^. In addition, the involvement of neutrophils in various pulmonary diseases is well-documented^40^. Thus, we expected to observe differences in macrophage and neutrophil levels in both sJIA-daPAP and non-sJIA-PAP. Importantly, eosinophils were the most enriched cell subset in LPRs across sJIA-daPAP and non-sJIA-PAP.

In sJIA-daPAP, the elevation of eosinophils can be linked to the abundance of serum TARC/CCL17, a chemokine that recruits eosinophils to the lung^41–43^. We have previously demonstrated that TARC/CCL17 serves as a serum marker for this PAP in sJIA, showing significant upregulation compared to levels noted in both sJIA without lung complications and sJIA with MAS^13^. Herein, the increased levels of TARC/CCL17 and eosinophils in sJIA-daPAP corroborate each other, highlighting their association as characteristics of drug-induced PAP in the context of sJIA. Taken together, enrichment of eosinophils and elevation of TARC/CCL17 can be considered as a feature of sJIA-daPAP.

Mouse models of MAS demonstrated pulmonary activation of IFN-γ pathways, which is postulated to link MAS biology with sJIA-associated lung disease^5,44^. Not surprisingly, peripheral blood from patients with sJIA commonly exhibits an IFN transcriptional signature^13,45^. In line with these results, we found that, unlike other inflammatory mediators, IFNγ levels are uniquely upregulated in lipoproteinosis regions of sJIA-daPAP, independent of concurrent MAS. IFNγ is primarily displayed in association with Th-1 cytokine signature, implicating inflammation and elevation of IFNγ is DReSS associated^46,47^. Interestingly, in sJIA-daPAP patients, our study reveals the derivation of IFNγ from granulocytes, particularly eosinophils, enriched within lipoproteinosis. Similarly, an increase in eosinophils-derived IFNγ has been reported in patients with aspirin-exacerbated respiratory disease (AERD) and differentiates AERD from drug-tolerant asthma^29^. Eosinophil-derived IFNγ secretion may represent a characteristic feature of DReSS-associated complications, including sJIA-daPAP.

Eosinophils are typically recruited to the tissue by type 2 cytokines and support Th-2 immune responses. Indeed, both purified mouse and human eosinophils can engage in Th-2 responses by secreting IFNγ, with these processes being heavily coordinated by cross-regulatory signals essential for Th-2 responses^48,49^. Given the prominent role of M2 macrophages in sJIA-daPAP, a high involvement of type 2 cytokines and Th-2 responses is anticipated in these cases. Combined, we hypothesize that sJIA-daPAP is mediated by the Th-2 signaling, with excessive IFNγ generation by eosinophils potentially serving as a co-mechanism that amplifies Th-1 pathways, thereby balancing Th-1/Th-2 immune responses. Additionally, eosinophil-derived IFNγ appears to have a dual role: at low concentrations, it can extend eosinophil survival; whereas at high concentrations, it can induce eosinophil apoptosis^29^. While overall eosinophil proportions do not differ between sJIA-daPAP and non-sJIA-PAP involved regions, this dual IFNγ function might explain the reduced eosinophil enrichment observed in sJIA-daPAP compared to non-sJIA-PAP lipoproteinosis regions, as higher IFNγ levels in sJIA-daPAP may compromise eosinophil numbers. Steroid treatment at the time of lung biopsy in our sJIA-daPAP cases and in one of two non-sJIA-PAP may also affect eosinophil enrichment in the tissues sampled.

IFNγ can elicit a response from eosinophils, including the induction of HLA-DR expression both *in vitro* and during trans-endothelial migration of eosinophils^31,50–54^. Besides IFNγ, HLA-DR on eosinophils is also inducible by GM-CSF, an immune modulator typically maintained within normal ranges in this drug associated PAP^5,54^. The pronounced elevation of HLA-DR expression on eosinophils in our sJIA-daPAP cases may result from the combined effects of IFNγ and GM-CSF regulation. Additionally, eosinophils’ HLA-DR expression can also be triggered by exogenous substances, such as segmental antigen in BAL^54,55^. Once HLA-DR is expressed by eosinophils, these cells are capable of processing antigens and presenting them to T cells, thereby amplifying the immune response^54^. We found that two of the five lung biopsies tested for HHV6 were positive for HHV6 in our sJIA-daPAP cohort. Although the temporal relationship between DReSS onset and HHV-6 reactivation remains controversial, herpes viral re-activations are part of DreSS, and evidence of HHV6 reactivation is an element in the Japanese consensus used to diagnose this type of severe delayed drug reaction^47^. Given the role of antigen-presenting molecules in immune regulation, we propose that the enriched eosinophils can enhance immune response and pathogen defense by expressing HLA-DR in response to elevated IFNγ expression. The significant presence of IFNγ-expressing eosinophils alongside the co-expression of HLA-DR can additionally serve as the signature of sJIA-daPAP.

MMP9 is a proteolytic enzyme involved in the breakdown of extracellular matrix components. In addition to being released by bronchial epithelial cells, type II alveolar cells, endothelial cells, and leukocytes, MMP9 can also be secreted from the tertiary granules of neutrophils. The significance of MMP9 on neutrophils is underscored by its critical role in neutrophil transmigration, with elevated MMP9 levels correlating with accelerated neutrophil infiltration into the lung^56–60^. Additionally, MMP9 is involved in the transmigration of eosinophils; therefore, the MMP9 we observe on neutrophils may explain and encourage eosinophilia in LPRs^61^. Our prior work identified MMP7, another matrix metalloproteinase, as a serum marker distinguishing sJIA with pulmonary complications developing during treatment with inhibitors of IL-1 and IL-6 from sJIA-MAS and active sJIA. The concurrent increases of MMP7 and MMP9 levels reflect an accelerated turnover rate of basement membranes, potentially facilitating the recruitment and entry of neutrophils and eosinophils into the inflamed lung tissue and may relate to neutrophilic and eosinophilic BAL found at the time of lung biopsy in sJIA-daPAP subjects. This appears concordant with previous reporting of neutrophilic BAL with negative bacterial cultures^3^ and contrasts with typically lymphocytic BAL in connective tissue associated pulmonary disease^62^ and a case report in autoimmune PAP^63^.

Overall, we present the first analysis of PAP leveraging a novel and high-throughput imaging platform to characterize the immune phenotypes associated with sJIA-daPAP.

MIBI-TOF analysis reinforces previously recognized lymphocytic infiltration in lung tissue and accumulation of innate immune cells within lipoproteinosis regions. Additionally, we propose the use of IFNγ-expressing eosinophils, with their co-expression of HLA-DR, as experimental indicators of sJIA-daPAP in children suspected of having this lung complication. Our results provide valuable insights into the possible mechanisms associated with this DReSS-related PAP occurring in sJIA patients and offer a resource for researchers investigating the pathogenesis of this serious condition. Since rheumatic diseases carry an increased risk of disease-associated lung complications, which in turn hinders diagnosing drug associated pulmonary complications^65,66^, future characterization of lung pathologies utilizing highly multiplexed imaging methods may simplify crucial diagnostic considerations.

### Limitations of the study

This descriptive study has several limitations. As the different forms of PAP are rare and given the invasiveness of biopsies, acquiring pulmonary specimens from pediatric patients is challenging. The small sample size of genetic mutation-associated PAP for non-sJIA-PAP cases does not fully represent the spectrum of non-drug associated PAP phenotypes; and the clinical heterogeneity among the sJIA-daPAP and non-sJIA-PAP categories can introduce confounding variables and impact the generalizability of the findings. sJIA-daPAP patients were treated with IL-1/IL-6 inhibitors combined with other immune suppressive medications for various durations prior to sampling and biopsy. It is difficult to isolate and account for treatment effects. We noted macrophage-derived multinucleated giant cells were seen only in non-sJIA-PAP samples and, while formation of these cells may depend on IL-6^64^, the limited comparisons presented here suggest further study. Finally, while IFNγ assays provide valuable insights into the cytokine signature of sJIA-daPAP, our study did not account for other common cytokines and chemokines. Future studies exploring various type 1 and type 2 cytokine profiles may provide more detail in the nuanced pathways involved in sJIA-daPAP.

## Methods

### Cohort Description and Region Selection

#### PAP Samples

All human samples were acquired in accordance with Stanford University’s institutional review board protocols (34679 and 13932) and as required at participating centers. Formalin-fixed paraffin-embedded (FFPE) pediatric PAP tissues (sJIA-daPAP: n = 12 from 9 patients; non-sJIA-PAP: n = 2 from 2 patients) were collected from the tissue repository of participating institutions. All clinical details for these specimens can be found in **Tables S1 and S2**. Serial sections (5 μm thickness) of each specimen were stained with hematoxylin and eosin (H&E) and inspected by an anatomic lung pathologist to screen for the presence of inflammation and lipoproteinosis. Three to six fields-of-view (FOVs), each measuring 500um x 500um, were chosen from each tissue block for imaging **(Figure 1a).** From four sJIA-daPAP specimens and one non-sJIA-PAP specimen, an adjacent uninvolved lung FOV was acquired. Visualization of FOV selection for one such specimen is shown in **Figure 1b**. No statistical methods were used to predetermine sample sizes.

#### Non-PAP and other staining control tissues

Pediatric controls consisted of uninvolved lung tissue, adjacent to a tumor, (n = 1 from one patient, termed “normal”) and lung tissue during pneumonia (n = 1 from one patient) from a tissue repository at a participating institution. All clinical details for these control specimens can be found in **Table S4**. These two samples are termed as “non-PAP” samples for the purpose of this study. Five 500 μm × 500 μm FOVs were chosen from both these tissue blocks for imaging. Additionally, control tissues from FFPE lymph node, tonsil, spleen, placenta, lung, colon, and liver were acquired from Stanford Health Care. Small regions of each tissue were selected and placed in a tissue microarray. One 500 μm × 500 μm FOV was chosen from each core for imaging. These tissues were used as staining controls, and representative FOVs for each marker are shown in **Figure S1**. H&E stains of non-PAP FOVs are included in **Figure S2**, and the MIBI-TOF images from non-PAP tissues were also used as technical control inputs in downstream analyses.

### Assay Preparation and MIBI-TOF Imaging

#### Slide preparation

Tissues were sectioned (5 μm section thickness) from tissue blocks on gold- and tantalum-sputtered microscope slides for MIBI-TOF imaging.

#### Antibody preparation

Antibodies were conjugated to isotopically-pure metal reporters as described previously^23,67^. Following conjugation, antibodies were diluted in Candor PBS Antibody Stabilization solution (Candor Bioscience, Cat# 130 050). The conjugated antibodies were either stored at 4 °C or lyophilized in 100 mM D-(+)-Trehalose dehydrate (Sigma-Aldrich) with ultrapure distilled H2O for storage at −20 °C. Before staining, lyophilized antibodies were reconstituted in a buffer of Tris (Thermo Fisher Scientific), sodium azide (Sigma-Aldrich), ultrapure water (Thermo Fisher Scientific) and antibody stabilizer (Candor Bioscience) to a concentration of 0.05 mg ml^-1^. The antibodies, metal reporters, and staining concentrations are listed in **Table S3.** Updated antibody preparation protocols are available^68^.

#### Tissue staining

MIBI-TOF slides were baked at 70 °C overnight, followed by deparaffinization and rehydration with washes in xylene (3×), 100% ethanol (2×), 95% ethanol (2×), 80% ethanol (1×), 70% ethanol (1×) and ddH2O with a Leica ST4020 Linear Stainer (Leica Biosystems). The antigens on the slides were retrieved by submerging slides in 3-in-1 Target Retrieval Solution (pH 9, DAKO Agilent, Cat# S2375) and incubating at 97 °C for 40 min in a Lab Vision PT Module (Thermo Fisher Scientific). After cooling to room temperature, slides were washed in 1× PBS IHC Washer Buffer with Tween 20 (Cell Marque), 0.1% (w/v) bovine serum albumin (Thermo Fisher Scientific), hereafter wash buffer. All tissues were blocked twice in blocking buffers at room temperature. The first block was for 30 min with endogenous biotin and avidin with an Avidin/Biotin Blocking Kit (BioLegend) and the second block was for 1 hr with 1× TBS IHC Wash Buffer with Tween 20 with 3% (v/v) normal donkey serum (Sigma-Aldrich), 0.1% (v/v) cold fish skin gelatin (Sigma-Aldrich), 0.1% (v/v) Triton X-100, and 0.05% (v/v) sodium azide. In between blocking steps, tissues were washed with wash buffer. The first antibody cocktail was prepared in 1x TBS IHC Wash Buffer with Tween 20 and 3% (v/v) normal donkey serum (Sigma-Aldrich) and filtered through a 0.1-μm centrifugal filter (Millipore) prior to incubation with tissue overnight at 4 °C in a humidity chamber. Following the overnight incubation, slides were washed twice for 5 min in wash buffer. The second day, an antibody cocktail was prepared as described above and incubated with the tissues for 1 hr at 4 °C in a humidity chamber. Following staining, slides were washed twice for 5 min in wash buffer and fixed in a solution of 2% glutaraldehyde (Electron Microscopy Sciences) solution in low-barium PBS for 5 min. Slides were washed in PBS (1×), 0.1 M Tris at pH 8.5 (3×) and ddH2O (2×) and then dehydrated by washing in 70% ethanol (1×), 80% ethanol (1×), 95% ethanol (2×) and 100% ethanol (2×). Slides were dried under vacuum prior to imaging. Updated tissue staining protocols are available^68,69^.

#### Imaging

Imaging was performed using a custom MIBI-TOF instrument with a Xe+ primary ion source, as described previously^70^. The imaging parameters were: acquisition setting, 100 kHz; field size, 500 μm x 500 μm at 1024 ×1024 pixels; dwell time, 2 ms; and ion dose, approximately 4.10 nAmp h per mm^2^. For each FOV, mass-spec pixel data were then converted to TIFF images where the counts for each mass were taken between the ‘Start’ and ‘Stop’ values defined in **Table S3**. Updated MIBI-TOF image acquisition protocols are available^68^.

### Data Analysis

#### Low-level processing

Multiplexed image sets were extracted; slide background-, adduct-, and oxide-subtracted, then denoised, and aggregate filtered as described previously^23,67^. There was evidence of non-specific signal in regions of lipoproteinosis. To remove the non-specific signal in FOVs exhibiting lipoproteinosis as determined by the expert pathologist, a lipoproteinosis subtraction mask was produced by subtracting calprotectin signal with a nuclear mask.

The nuclear mask was formed by merging the expressions of H3K9ac and H3K27me3, followed by applying a Gaussian filter, applying a threshold, and implementing a 5-pixel radial expansion. Finally, the resulting masks were subtracted from all marker layers to eliminate the lipoproteinosis-sticking signal. The resulting pre-processed images were Gaussian blurred using a standard deviation of two. Pixels were normalized such that the total expression of each pixel was equal to one. A 99.9% normalization was applied for each marker.

#### Single-cell segmentation

Cell segmentation was performed using the pre-trained Mesmer^24^ convolutional neural network architecture with a combination of HH3, H3K9ac, and H3K27me3 as the nuclear channel and CD45, panCK, and Na+/K+ ATPase as the cell membrane channel. All PAP and non-PAP FOVs were provided as input to the network to predict the class of each pixel: nuclear interior, nuclear border, or non-nuclear background. The nuclear interior probability map for each image was thresholded and segmented, followed by a 3-pixel radial expansion around each nucleus to define the cell object boundaries. A correction was applied to FOVs that contained multinucleated giant cells (giant cells). Each giant cell was identified using a combination of HH3, CD45, and vimentin and manually segmented in ImageJ to produce a binary mask of each giant cell. All pixels within the binary mask were reassigned to belong to the giant cell object(s).

#### Single-cell phenotyping and composition

MIBI-TOF FOVs from PAP and non-PAP tissues were included as inputs for single-cell phenotyping, with non-PAP tissues included as technical controls. Employing Pixie^25^, pixels were clustered into 100 clusters using a self-organizing map (SOM) based on the expression of 23 phenotypic markers: CD4, CD14, Foxp3, CD31, EPOX, CD209, CD206, iNOS, CD68, CD11c, CD8, CD3, CD163, CD20, CD16, HLA-DR, CD57, CD45, panCK, calprotectin, vimentin, SMA, and tryptase. The average expression of each cluster was found, and the z-score for each marker across the 100 pixel clusters was computed and capped at three. Using z-scored expression values, the pixel clusters were grouped into 20 meta clusters using consensus hierarchical clustering. Next, by applying the segmentation border masks, we counted the number of each of the 20 pixel meta clusters in each cell. These counts were normalized by cell size. Using these frequency measurements as the feature vector, the cells were clustered using a SOM into 100 cell clusters. Similar to the pixel clusters, the average expression of each of these clusters was found, and the z-score was computed with a maximum z-score of three. These clusters were grouped using consensus hierarchical clustering into 20 cell meta clusters. By assessing a heatmap of the 100 cell clusters, a few of the clusters were manually reassigned to the appropriate meta cluster. Each of the cell meta clusters was then manually annotated with its cell phenotype by assessing marker expression. Giant cells were separated into their own meta cluster. Cell phenotype maps (CPMs) for all PAP and non-PAP sample FOVs are shown in **Figure S2**.

#### Differential Abundance Analysis

To perform differential abundance analysis, the diffcyt R package^71^ statistical framework was used. Specifically, generalized linear mixed models were employed to test for differences in cluster abundance and cluster marker expression across conditions. For each cluster, we fit a binomial regression model in which we model the log-odds (and thus the proportion of cells 𝑝*_ij_*) of each cluster in a given patient *i* and a given FOV *j* according to the following equation:

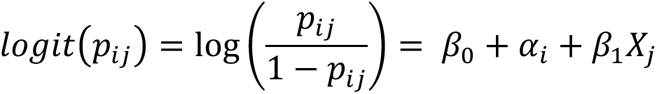

where 𝑝*_ij_* gives the proportion of cells in a given cluster in patient *i* and FOV *j*; 𝛼*_i_* gives the random intercept for each patient *i* (in which 𝛼*_i_* ∼ 𝑁(0, 𝜎) and σ is estimated during model fitting); 𝑋*_j_* is an indicator variable representing whether or not a FOV *j* was taken from a given experimental condition (1 if yes; 0 otherwise), and all β’s are model parameters optimized during model fitting. Using this setup, we can apply null-hypothesis significance testing to 𝛽_1_: if 𝛽_1_ is significantly different from 0 in the model, the proportion of cells in a given cluster differs significantly between the levels of the experimental variable, while controlling for individual-to-individual variation.

#### Lipoproteinosis, Mask Generation and Corresponding Analysis

Using the lipoproteinosis subtraction mask (described above), for each FOV, a rudimentary lipoproteinosis mask (**Figure S2**) was refined via manual annotation with Adobe Photoshop based on vimentin expression and visualization of the corresponding hematoxylin and eosin-stained tissue regions. The binary masks were confirmed by the expert pathologist. Cells were deemed lipoproteinosis+ if their centroid was located in a region of positivity in the lipoproteinosis mask.

#### Enrichment Analysis

To quantify the enrichment of certain cell types within lipoproteinosis pathology, for each cell type, the number of lipoproteinosis+ and lipoproteinosis-cells were determined on a per-FOV basis. Their proportion was then calculated out of the total number of lipoproteinosis+ and lipoproteinosis-cells, respectively, per FOV. Finally, for each cell type, the log2 of the ratio of the lipoproteinosis+ cell proportion and the lipoproteinosis-cell proportion was determined. In this analysis, the relationship of enrichment is dependent on the fact that there are both lipoproteinosis+ and lipoproteinosis-cells for every cell type in each FOV. To measure complete observations, partial observations were dropped. The same analysis was applied for both cell phenotypes and cell microenvironments.

#### Differential Expression Analysis

Comparing marker expression between lipoproteinosis+ cells and lipoproteinosis-cells, a two-tailed Student’s t-test was performed for every functional marker (CD45RO, CD57, GrzB, H3K27me3, H3K9ac, HLA-DR, HO-1, IDO-1, IFNγ, iNOS, Ki67, MMP9, pS6, and TIM-3) in each FOV. The corresponding p-values were corrected using the Benjamini/Hochberg correction. For the expression of each functional marker, the log2 fold change between lipoproteinosis+ and lipoproteinosis-cells was also calculated. The results of the differential expression analysis were reported in a volcano plot. Unless otherwise stated, we used a significance threshold of absolute value of log2FC > 1 and adjusted p-value of 0.05.

#### UMAP Visualization

UMAP^72^ embeddings were determined for all eosinophils, neutrophils, and M2 macrophages using the python implementation with the parameters n neighbors = 15 and min_dist = 0.5 and all markers: CD11c, CD14, CD16, CD163, CD20, CD206, CD209, CD3, CD31, CD4, CD45, CD45RO, CD57, CD68, CD8, calprotectin, EPOX, Foxp3, GrzB, HH3, H3K27me3, H3K9ac, HLA-DR, HO-1, IDO-1, IFNγ, Ki67, MMP9, NaKATPase, panCK, SMA, TIM-3, tryptase, vimentin, iNOS, and pS6.

#### Spatial-Latent Dirichlet Allocation (spatial-LDA)

Spatial-LDA^34^ was conducted to identify microenvironments (MEs) across all FOVs from PAP and non-PAP tissues; non-PAP tissue FOVs were included as technical controls. A spatial radius of r = 50 μm (100 pixels) and a ME number of nine were used. The ME number was determined empirically. For each FOV, a maximum probability map was produced by assigning each cell to the ME with the highest probability and coloring that cell by its ME. ME maps for all PAP and non-PAP sample FOVs are shown in **Figure S2**.

#### Software and Reproducibility

Data collection and analysis were not performed blind to disease category. Image processing was conducted with MAUI (https://github.com/angelolab/MAUI) in MATLAB 2019b. Cell segmentation was performed with Mesmer (https://www.deepcell.org/predict). Pixel clustering and cell phenotyping were conducted with Pixie (https://github.com/angelolab/pixie). The Google Colab notebook (https://colab.research.google.com/drive/1Dx8nW37OFaRMN6ILfrwQ6nhDAMQAdNH4?usp=sharing) from the spatial_lda GitHub repository was used for implementation of spatial-LDA. Statistical analysis was conducted in MATLAB 2019b, R v4.4.1, and Python 3.11.4. Data visualization and plots were generated in R and Python. Representative images were processed in Adobe Photoshop; processed images were displayed to show representative imaging data, but quantification of results was performed on unaltered images. Figures were prepared in Adobe Illustrator. Schematic visualizations were produced at https://biorender.com.

### Resource Availability

#### Lead contact

Further information and requests for resources and reagents should be directed to and will be fulfilled by the lead contact, Erin F. McCaffrey.

#### Materials availability

This study did not generate new unique reagents.

#### Data and code availability

All processed images, cell segmentation masks, lipoproteinosis masks, annotated single-cell data, and per-FOV meta data are deposited in Mendeley’s data repository and can be accessed using the following link: https://doi.org/10.17632/pbywrb297x.1.

All custom code used to analyze data has been deposited in GitHub and can be accessed using the following link: https://github.com/angelolab/publications/tree/main/2024-Delmastro_etal_sJIA-daPAP.

## Acknowledgements

We thank Pauline Chu and the Stanford Human Histology Core for providing technical assistance. We acknowledge Scott W Aesif. MD, PhD and Abdur R Khan, MD, who assisted in providing specimens for this study and Grant Schulert, MD, PhD, who assisted in providing clinical details. I.A. is an awardee of the Weizmann Institute of Science-Israel Women’s Postdoctoral Career Development Award in Science. M.A. is supported by 5U54CA20997105, 5DP5OD01982205, 1R01CA24063801A1, 5R01AG06827902, 5UH3CA24663303, 5R01CA22952904, 1U24CA22430901, 5R01AG05791504 and 5R01AG05628705 from the NIH, W81XWH2110143 from the DOD, and other funding from the Bill and Melinda Gates Foundation, the Cancer Research Institute, the Parker Center for Cancer Immunotherapy, the Lucile Packard Foundation for Children’s Health, and the Breast Cancer Research Foundation. E.F.M. is supported by the Division of Intramural Research, NIAID/NIH.

## Author contributions

Conceptualization, E.M.; Software, A.D., E.F.M., and T.J.K.; Formal Analysis, A.D., E.F.M., C.C.L., T.J.K.; Investigation, A.D. and E.F.M.; Resources, S.Y.T., G.D., and E.M.; Data Curation, A.D., E.F.M, I.A., and M.B.; Writing - Original Draft, A.D.; writing – Review & Editing, A.D., E.F.M., X.D., V.E.S, and S.S.; Supervision, E.F.M., M.A., V.E.S., and E.M.

