## Supplemental Tables and Figures for "IFNγ-Expressing Myeloid Cells Localize within Lipoproteinosis during Drug-Associated Pulmonary Alveolar Proteinosis occurring in Systemic Juvenile Idiopathic Arthritis"

### Supplemental Figures

| Patient ID | 1 | 2 | 3 | 4 | 5 | 6 | 7 | 8 | 9 |
|---|---|---|---|---|---|---|---|---|---|
| **Baseline demographics** | | | | | | | | | |
| Primary illness | sJIA | sJIA-like | sJIA | sJIA | sJIA-like | sJIA-like | sJIA | sJIA | sJIA |
| Age at onset of primary illness (years) | 1.1 | 1.4 | 8.8 | 9.9 | 1.5 | 0.6 | 1.1 | 0.9 | 5.8 |
| Sex | F | M | F | M | F | F | M | M | M |
| Self-identified ethnicity | White | White | Middle Eastern | White | Hispanic | White | White/SE Asian | White | White |
| Other comorbidity | none | severe BPD | none | T21/CHD | none | T21/CHD | T21 bal. transl. | none | none |
| HLA-DRB1* | 15*/15* | 15*/X | 13*/11* | 15*/15* | 15*/11* | 15*/12* | 15*/12* | 15*/X | 15*/15* |
| **Inhibitors of Interleukin-1/Interleukin-6** | | | | | | | | | |
| Age at first exposure to IL-1i/IL-6i (years) | 1.1 | 1.6 | 9 | 10 | 4 | 0.6 | 1.1 | 1.1 | 5.8 |
| First IL-1i/IL-6i used as treatment | anakinra | anakinra | canakinumab | tocilizumab | tocilizumab | anakinra | anakinra | anakinra | tocilizumab |
| Subsequent IL-1i/IL-6i before lung biopsy | canakinumab | canakinumab | tocilizumab | canakinumab | canakinumab | none | tocilizumab | canakinumab | anakinra & canakinumab |
| RegiSCAR-DReSS implicating IL-1i/IL-6i | definite | definite | definite | definite | definite | definite | definite | definite | definite |
| **At lung biopsy date** | | | | | | | | | |
| Age (years) | 1.6 | 2 | 9.8 | 11 | 6 | 2 | 3 | 3 | 8 |
| IL-1i/IL-6i treatment | anakinra | canakinumab | anakinra | canakinumab | canakinumab | anakinra | canakinumab | tocilizumab | anakinra & canakinumab |
| cumulative duration of IL-1i/IL-6i | 6 months | 7 months | 9 months | 1.6 years | 2 years | 2 years | 2.2 years | 2.2 years | 2.7 years |
| Prednisone equivalent dose (mg/kg/day) | 0.9 | 1 | 1 | 0.9 | 0.6 | 0.8 | 0.4 | monthly IV only | 0.2 plus IV pulse |
| Other immune suppressive medications | MMF | none | CSA | methotrexate | cyclosporine | cyclosporine | MMF | methotrexate | MMF |
| Clinical data (within 1 month of biopsy) | | | | | | | | | |
| MAS | yes | no | no | no | no | no | no | no | no |
| Serum ferritin | 1500 | 338 | 11,000 | 451 | 28 | 102 | 271 | 49 | 247 |
| C-reactive protein (CRP) | 5.8 mg/dl | 1.3 mg/dl | 1.5 mg/dl | 2.05 mg/dl | below range | below range | not done | below range | 6.1 mg/dl |
| Active arthritis | no | no | no | no | yes | no | no | no | no |
| BAL nucleated cell counts† | 42M/46N/3L/9E | 59M/36N/2L/0E | 97M/2N/0L/1E | 72M/18N/10L/0E | unavailable | unavailable | 65M/21N/14L/0E | 82M/6N/12L/0E | 72M/23N/3L/1E |
| HHV6-PCR result in lung biopsy tissue | not tested | not tested | not tested | positive | positive | negative | not tested | negative | negative |
| **Participation in previous cohorts** | | | | | | | | | |
| Included in publications 1, 2, 3, 4, or 5 | 2,4 | 4 | 2,4 | 1,2 | none | 1,2 | 1-3 | 2,4 | 1-3 |
| **Per patient feature visualization** | | | | | | | | | |

Cell Subset Proportions

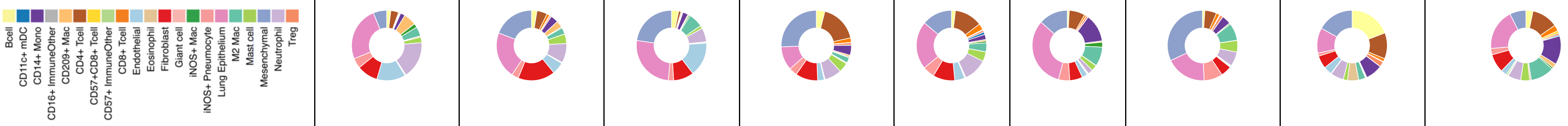


Microenvironment Proportions

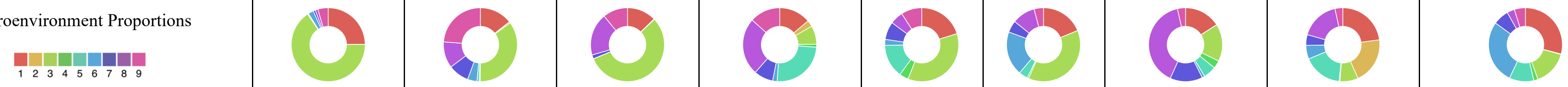


sJIA, systemic juvenile idiopathic arthritis per ILAR[42]; sJIA-like, similar illness failing to classify as sJIA yet treated similarly; severe BPD, oxygen dependent bronchopulmonary dysplasia of prematurity; T21, Trisomy 21; T21 bal. transl., balanced translocation of Trisomy 21 with no clinical effect; CHD, severe congenital heart disease requiring neonatal surgical repair; HLA-DRB1*15/X, presumed homozygous DRB1*15; IL-1i/IL-6i, inhibitor of IL-1 (anakinra or canakinumab) or IL-6 (tocilizumab); RegiSCAR-DReSS, score ≥6 is "definite" drug reaction with eosinophilia and systemic symptoms (DReSS) per validated RegiSCAR scoring[43] (2); MAS, macrophage activation syndrome per Ravelli classification[44]; BAL, bronchoalveolar lavage fluid; MMF, mycophenolate mofetil; CRP, C-reactive protein; < range, result lower than the laboratory's reporting range; HHV6, human herpes virus 6; Publications 1, 2, 3, 4, or 5, 1=Saper et al 2019[3], 2= Saper and Ombrello et al 2021[10], 3= Chen et al 2022[12], 4= Saper et al 2024[9], 5= Neehus et al 2024[28]

†normal range of nucleated cell counts in BAL per American Thoracic Society[45] (8): monocytes (M)>85%, lymphocytes (L)10-15%, neutrophils (N) ≤3%, eosinophils (E) ≤1%

**Table S1: sJIA-daPAP cohort clinical characteristics with per patient feature visualization.**

| Patient ID | 10 | 11 |
|---|---|---|
| Baseline demographics | | |
| Primary illness | GATA2/Fevers, weight loss | COPA/Diffuse alveolar hemorrhage |
| Age at onset of primary illness (years) | 18 | 1.8 |
| Sex | F | F |
| Self-identified ethnicity | White | Declined |
| Other comorbidity | Restrictive lung disease | PH, kidney disease |
| HLA-DRB1* | not determined | not determined |
| Inhibitors of Interleukin-1/Interleukin-6 | | |
| Age at first exposure to IL-1i/IL-6i (years) | N/A | N/A |
| First IL-1i/IL-6i used as treatment | N/A | N/A |
| Subsequent IL-1i/IL-6i before lung biopsy | N/A | N/A |
| RegiSCAR-DReSS implicating IL-1i/IL-6i | N/A | N/A |
| At lung biopsy date | | |
| Age (years) | 20.2 | 2.3 |
| IL-1i/IL-6i treatment | none | none |
| cumulative duration of IL-1i/IL-6i | N/A | N/A |
| Prednisone equivalent dose (mg/kg/day) | none | 1.0 |
| Other immune suppressive medications | none | Rituximab & cyclophosmamide |
| Clinical data (within 1 month of biopsy) | | |
| MAS | no | no |
| Serum ferritin | 177 | 2000 |
| C-reactive protein (CRP) | 8.2 mg/dl | 2.9 mg/dl |
| Active arthritis | no | no |
| BAL nucleated cell counts† | not determined | not determined |
| HHV6-PCR result in lung biopsy tissue | not tested | not tested |
| Participation in previous cohorts | | |
| Included in publications 1, 2, 3, 4, or 5 | 3,5 | none |
| Per patient feature visualization | | |
| Cell Subset Proportions 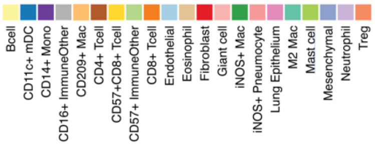  | 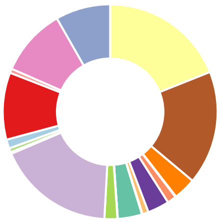 | 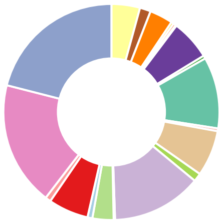 |
| Microenvironment Proportions 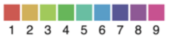  | 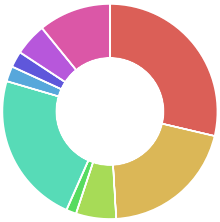 | 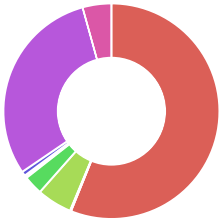 |

PH, pulmonary hypertension; HLA-DRB1*15/X, presumed homozygous DRB1*15; IL-1i/IL-6i, inhibitor of IL-1 (anakinra or canakinumab) or IL-6 (tocilizumab); RegiSCAR-DReSS, score ≥6 is "definite" drug reaction with eosinophilia and systemic symptoms (DReSS) per validated RegiSCAR scoring[43] (2); MAS, macrophage activation syndrome per Ravelli classification[44]; BAL, bronchoalveolar lavage fluid; MMF, mycophenolate mofetil; CRP, C-reactive protein; < range, result lower than the laboratory's reporting range; HHV6, human herpes virus 6; Publications 1, 2, 3, 4, or 5, 1=Saper et al 2019[3], 2= Saper and Ombrello et al 2021[10], 3= Chen et al 2022[12], 4= Saper et al 2024[9], 5= Neehus et al 2024[28]

†normal range of nucleated cell counts in BAL per American Thoracic Society[45] (8): monocytes (M)>85%, lymphocytes (L)10-15%, neutrophils (N) ≤3%, eosinophils (E) ≤1%

**Table S2: non-sJIA-PAP cohort clinical characteristics with per patient feature visualization.**

| Day 1 (overnight stain) | | | | | | | | Parameters for Analysis | | |
|---|---|---|---|---|---|---|---|---|---|---|
| **Antibody Target** | **Biological Relevance** | **Provider** | **Catalog Number** | **Lot** | **Clone** | **Mass Channel** | **Titer (µg/mL)** | **Start** | **Stop** | **AggFilter** |
| CD45RO | T cell memory | Biolegend | 304202 | B232536 | UCHL1 | 141Pr | 0.75 | 140.7 | 141 | 150 |
| HO-1 | Immunoregulation | Abcam | ab13248 | GR295624-7 | HO-1-1 | 142Nd | 0.25 | 141.7 | 142 | 200 |
| CD4 | T cells | Abcam | ab181724 | GR3215375-1 | EPR6855 | 143Nd | 0.25 | 142.7 | 143 | 200 |
| CD14 | Monocytes/Macrophages | Cell Signaling Technology | 56082BF | 2 | D7A2T | 144Nd | 0.25 | 143.7 | 144 | 200 |
| Foxp3 | Regulatory T cells (Tregs) | BD Biosciences | 624084 | 8099783 | 236A/E7 | 146Nd | 1.00 | 145.7 | 146 | 150 |
| CD31 | Endothelium | Abcam | ab212712 | GR3221684-8 | JC/70A | 148Nd | 0.25 | 147.7 | 148 | 150 |
| EPOX | Eosinophils | Custom antibody | NA | NA | NA | 149Sm | 0.75 | 148.7 | 149 | 200 |
| GrzB | Cytotoxic cells | Cell Signaling Technology | 46890BF | 2 | D6E9W | 150Nd | 0.50 | 149.7 | 150 | 100 |
| Ki67 | Proliferation | Cell Signaling Technology | 9449BF | 7 | 8D5 | 151Eu | 0.25 | 150.7 | 151 | 125 |
| CD209 | Myeloid-lineage cells | BD Biosciences | 624084 | 9178683 | DCN46 | 152Sm | 0.25 | 151.7 | 152 | 100 |
| CD206 | Anti-inflammaotry (M2-like) Macrophages | R&D Systems | MAB25341 | CEZR0217061 | 685645 | 153Eu | 0.25 | 152.7 | 153 | 200 |
| pS6 | Activation | Cell Signaling Technology | 4858BF | 12 | D57.2.2E | 154Sm | 0.25 | 153.7 | 154 | 100 |
| iNOS | Macrophage inflammation (M1-like) | Spring Bioscience | M4264 | 170802 | SP126 | 155Gd | 1.00 | 154.7 | 155 | 100 |
| CD68 | Macrophages | Cell Signaling Technology | 76437BF | 2 | D4B9C | 156Gd | 0.25 | 155.7 | 156 | 100 |
| CD11c | Myeloid-lineage cells | Abcam | ab216655 | GR32103491 | EP1347Y | 157Gd | 0.50 | 156.7 | 157 | 100 |
| CD8 | T cells | Cell Marque | 108M-OEM1404 | 1514101 | D8A8Y | 158Gd | 0.25 | 157.7 | 158 | 100 |
| CD3ε | T cells | Cell Signaling Technology | 85061BF | 4 | D7A6E | 159Tb | 0.25 | 158.7 | 159 | 100 |
| IDO1 | Immunoregulation | Abcam | ab55305 | GR239251-1 | ab55305 | 160Gd | 1.00 | 159.7 | 160 | 100 |
| TIM-3 | Immunoregulation | Abcam | ab242080 | GR32486051 | EPR22241 | 162Dy | 0.50 | 161.7 | 162 | 100 |
| CD163 | Anti-inflammaotry (M2-like) Macrophages | Cell Signaling Technology | 93498BF | 2 | D6U1J | 163Dy | 1.00 | 162.7 | 163 | 200 |
| CD20 | B cells | Cell Marque | 120M-8-OEm | 1908402 | L26 | 164Dy | 0.25 | 163.7 | 164 | 100 |
| CD16 | FcγRIII, Myeloid cells and Natural Killer cells | Cell Signaling Technology | 24326BF | 2 | D1N9L | 165Ho | 0.50 | 164.7 | 165 | 100 |
| IFNγ | Inflammatory cytokine | Abcam | ab218890 | GR3191590-2 | IFNG/466 | 166Er | 1.00 | 165.7 | 166 | 100 |
| HLA-DR | MHC class II | Abcam | ab215985 | GR3218615-1 | EPR3692 | 167Er | 0.25 | 166.7 | 167 | 100 |
| CD57 | Lymphocyte maturation | Abcam | ab212408 | GR3218433-1 | NK/804 | 168Er | 0.25 | 167.7 | 168 | 100 |
| CD45 | Immune cells | Cell Signaling Technology | 13917BF | 2 | D9M8I | 169Tm | 0.25 | 168.7 | 169 | 100 |
| H3K9Ac | Chromatin accessibility | Cell Signaling Technology | 9649BF | 12 | C5B11 | 170Yb | 0.50 | 169.7 | 170 | 150 |
| PanCK | Epithelium | ThermoFisher | MS-343-PABX | 19.2FZ343X1810A | AE1/AE3 | 171Yb | 0.25 | 170.7 | 171 | 100 |
| H3K27me3 | Chromatin accessibility | Cell Signaling Technology | 9733BF | 11 | C36B11 | 172Yb | 0.25 | 171.7 | 172 | 100 |
| MMP9 | Immunoregulation | Abcam | ab204850 | GR280040-3 | EP1254 | 173Yb | 0.25 | 172.7 | 173 | 100 |
| Calprotectin | Neutrophils | ThermoFisher | MA1-81381 | UE2778733 | MAC387 | 174Yb | 0.50 | 173.7 | 174 | 100 |
| NaKATPase | Cell membrane | Abcam | ab167390 | GR3229163-1 | EP1845Y | 175Lu | 0.50 | 174.7 | 175 | 100 |

| Day 2 (1 hour) | | | | | | | | Parameters for Analysis | | |
|---|---|---|---|---|---|---|---|---|---|---|
| **Antibody Target** | | **Provider** | **Catalog Number** | **Lot** | **Clone** | **Mass Channel** | **Titer (µg/mL)** | **Start** | **Stop** | **AggFilter** |
| HH3 | Nuclei | Cell Signaling Technology | 4499BF | 15 | D1H2 | 89Y | 2.00 | 88.7 | 89 | 150 |
| Vimentin | Cytoplasm | Cell Signaling Technology | 5741BF | 4 | D21H3 | 113In | 2.00 | 112.7 | 113 | 150 |
| SMA | Myofibroblasts | Spring | M4714.C | 150410 | SP171 | 115In | 2.00 | 114.7 | 115 | 200 |
| biotin | NA | Biolegend | 409002 | B267484 | 1D4-C5 | 149Sm | 0.50 | 148.7 | 149 | 200 |
| Tryptase | Mast cells | Abcam | ab216451 | GR31363111 | EPR9522 | 176Yb | 2.00 | 175.7 | 176 | 100 |

**Table S3: Multiplexed imaging antibody panel staining conditions and low-level processing parameters.**

| Patient ID | Patient 12 | Patient 13 |
|---|---|---|
| Baseline demographics | | |
| Primary illness | Rhabdomyosarcoma | Pneumonia |
| Age at onset of primary illness (years) | 14 | 2 |
| Sex | M | F |
| Self-identified ethnicity | White | White |
| Other comorbidity | none | Asthma |
| HLA-DRB1* | not determined | not determined |
| Inhibitors of Interleukin-1/Interleukin-6 | | |
| Age at first exposure to IL-1i/IL-6i (years) | N/A | N/A |
| First IL-1i/IL-6i used as treatment | N/A | N/A |
| Subsequent IL-1i/IL-6i before lung biopsy | N/A | N/A |
| RegiSCAR-DReSS implicating IL-1i/IL-6i | N/A | N/A |
| At lung biopsy date | | |
| Age (years) | 14.3 | 2.1 |
| IL-1i/IL-6i treatment | none | none |
| cumulative duration of IL-1i/IL-6i | N/A | N/A |
| Prednisone equivalent dose (mg/kg/day) | none | none |
| Other immune suppressive medications | none | none |
| Clinical data (within 1 month of biopsy) | | |
| MAS | no | no |
| Serum ferritin | not determined | not determined |
| C-reactive protein (CRP) | not determined | not determined |
| Active arthritis | no | no |
| BAL nucleated cell counts† | not determined | not determined |
| HHV6-PCR result in lung biopsy tissue | not tested | not tested |
| Participation in previous cohorts | | |
| Included in publications 1, 2, 3, 4, or 5 | none | none |
| Per patient feature visualization | | |
| Cell Subset Proportions 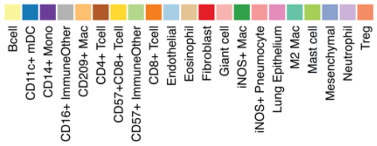  | 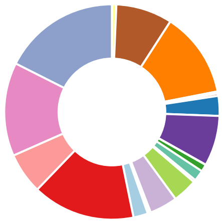 | 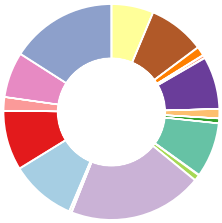 |
| Microenvironment Proportions 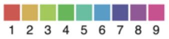  | 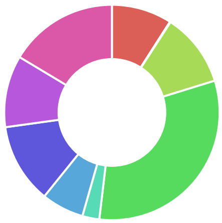 | 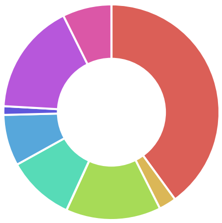 |

IL-1i/IL-6i, inhibitor of IL-1 (anakinra or canakinumab) or IL-6 (tocilizumab); RegiSCAR-DReSS, score ≥6 is "definite" drug reaction with eosinophilia and systemic symptoms (DReSS) per validated RegiSCAR scoring[43] (2); MAS, macrophage activation syndrome per Ravelli classification[44]; BAL, bronchoalveolar lavage fluid; MMF, mycophenolate mofetil; CRP, C-reactive protein; < range, result lower than the laboratory's reporting range; HHV6, human herpes virus 6; Publications 1, 2, 3, 4, or 5, 1=Saper et al 2019[3], 2= Saper and Ombrello et al 2021[10], 3= Chen et al 2022[12], 4= Saper et al 2024[9], 5= Neehus et al 2024[28]

†normal range of nucleated cell counts in BAL per American Thoracic Society[45] (8): monocytes (M)>85%, lymphocytes (L)10-15%, neutrophils (N) ≤3%, eosinophils (E) ≤1%

**Table S4: Pediatric control clinical characteristics with per patient feature visualization.**

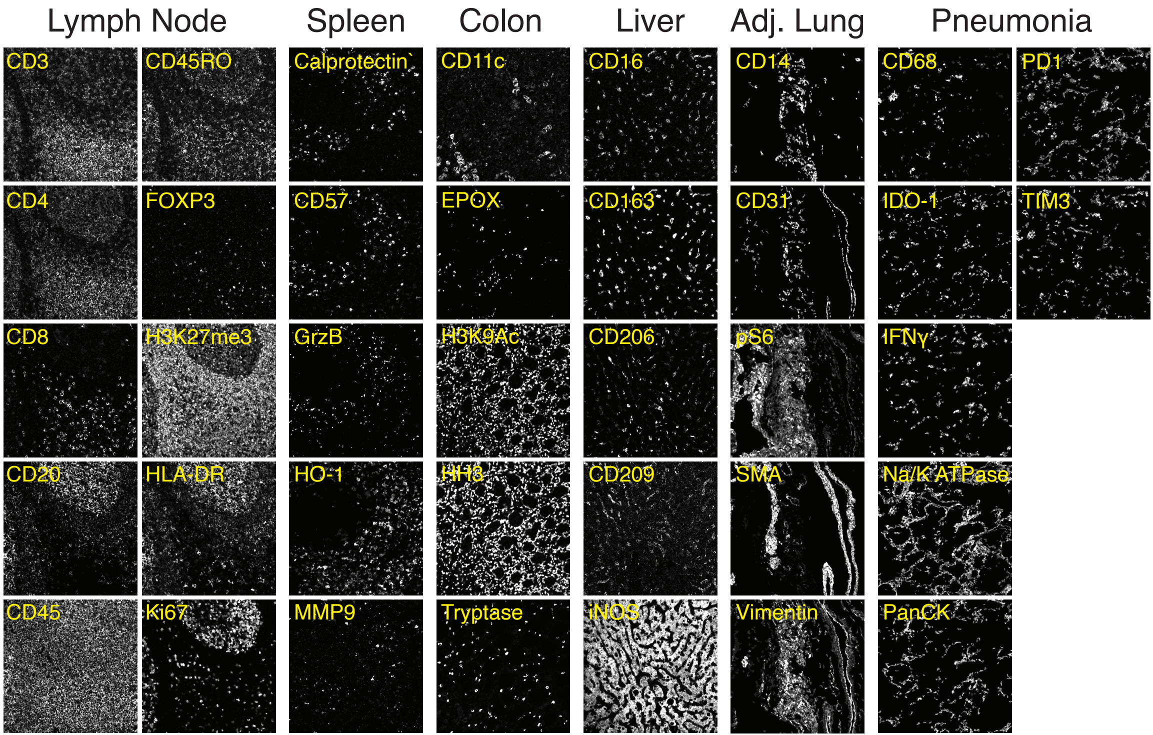

Lymph Node
Spleen
Colon
Liver
Adj. Lung
Pneumonia
CD3
CD45RO
Calprotectin`
CD11c
CD16
CD14
CD68
PD1
CD4
FOXP3
CD57
EPOX
CD163
CD31
IDO-1
TIM3
CD8
H3K27me3
GrzB
H3K9Ac
CD206
pS6
IFNγ
CD20
HLA-DR
HO-1
HH3
CD209
SMA
Na/K ATPase
CD45
Ki67
MMP9
Tryptase
iNOS
Vimentin
PanCK

**Figure S1: Multiplexed imaging staining controls.** Grayscale images of protein expression in control tissues (lymph node, spleen, colon, liver, normal lung adjacent to a tumor, and lung tissue from a pediatric pneumonia patient.)

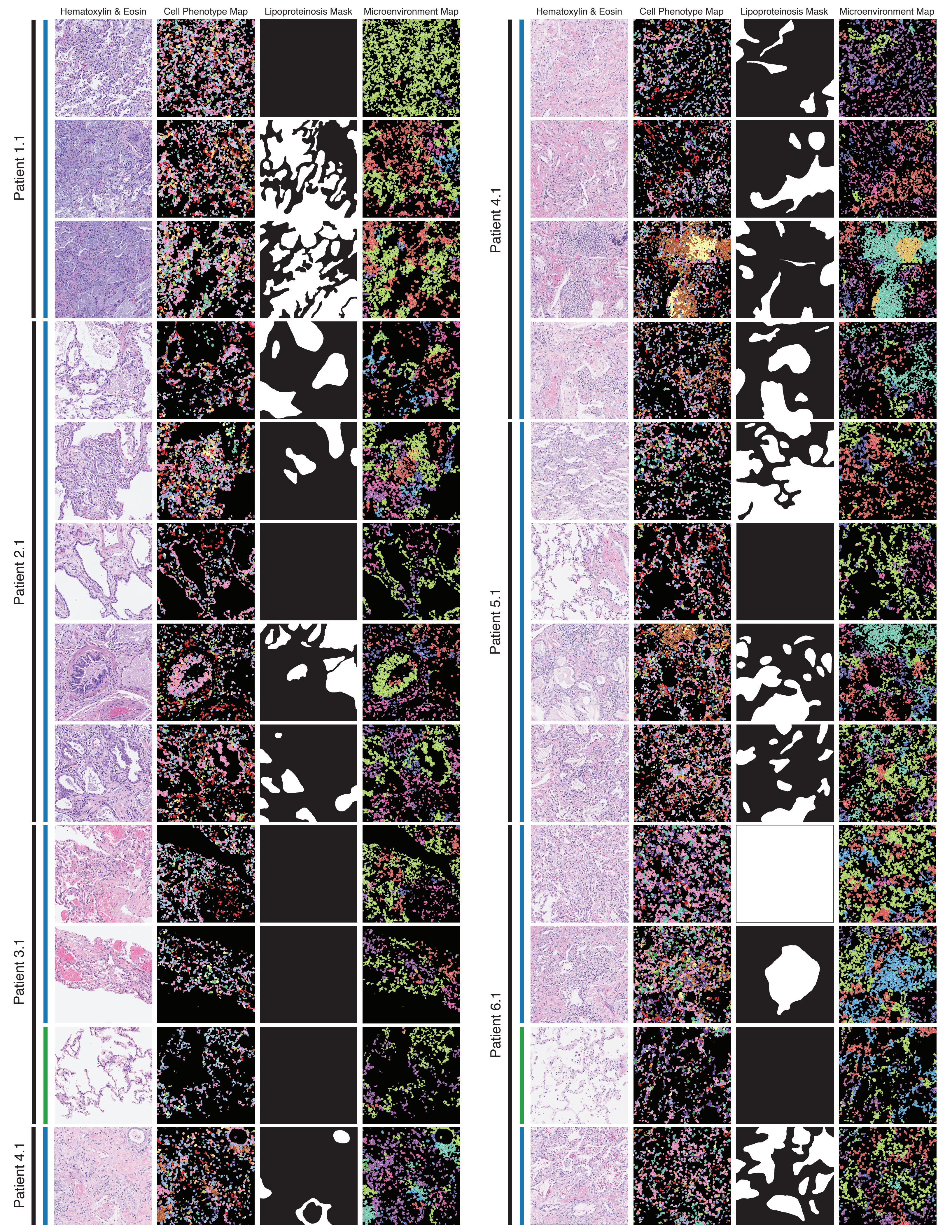
Hematoxylin & Eosin
Cell Phenotype Map
Lipoproteinosis Mask
Microenvironment Map
Patient 1.1
Patient 2.1
Patient 3.1
Patient 4.1
Patient 5.1
Patient 6.1

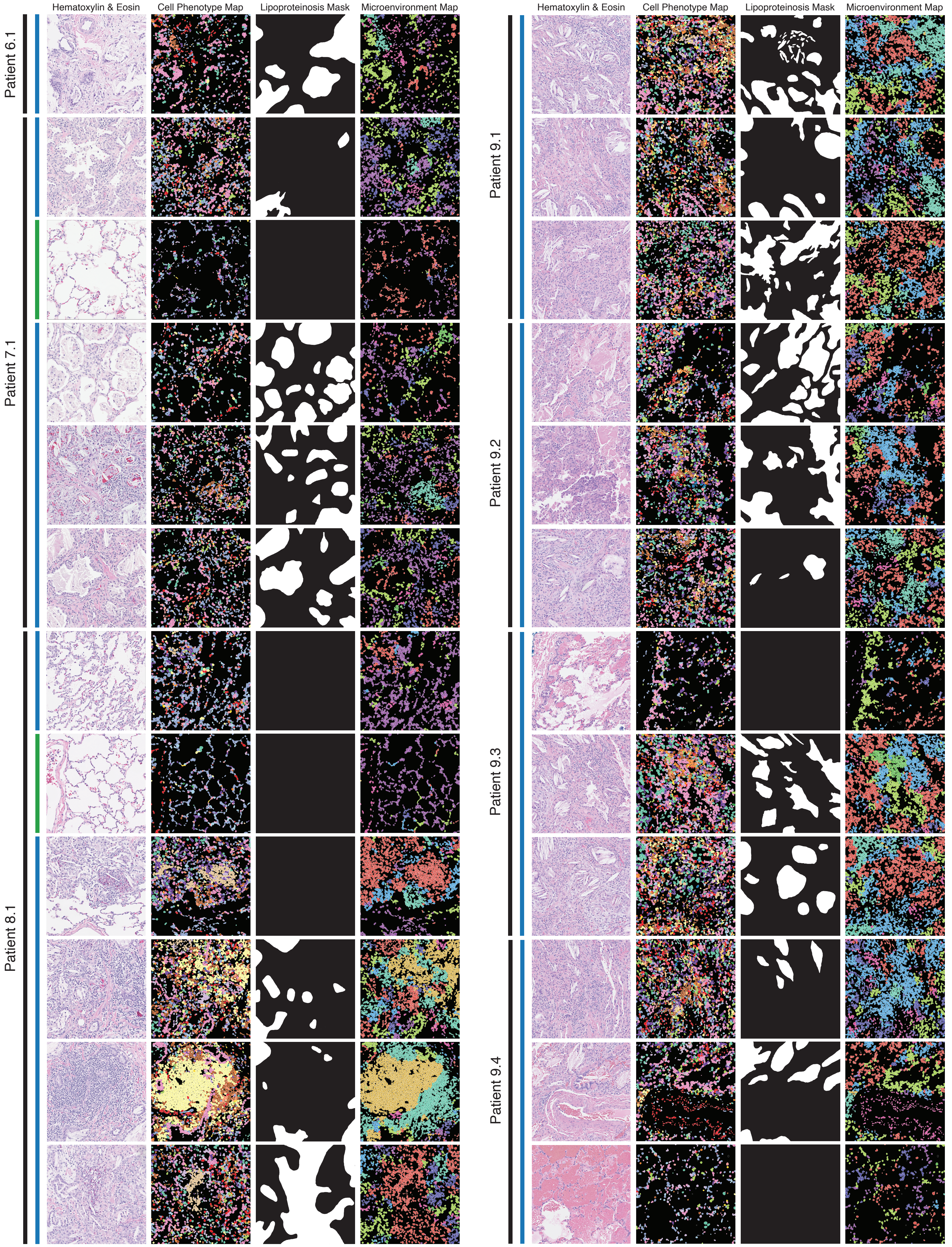

Hematoxylin & Eosin
Cell Phenotype Map
Lipoproteinosis Mask
Microenvironment Map
Patient 6.1
Patient 7.1
Patient 8.1
Hematoxylin & Eosin
Cell Phenotype Map
Lipoproteinosis Mask
Microenvironment Map
Patient 9.1
Patient 9.2
Patient 9.3
Patient 9.4

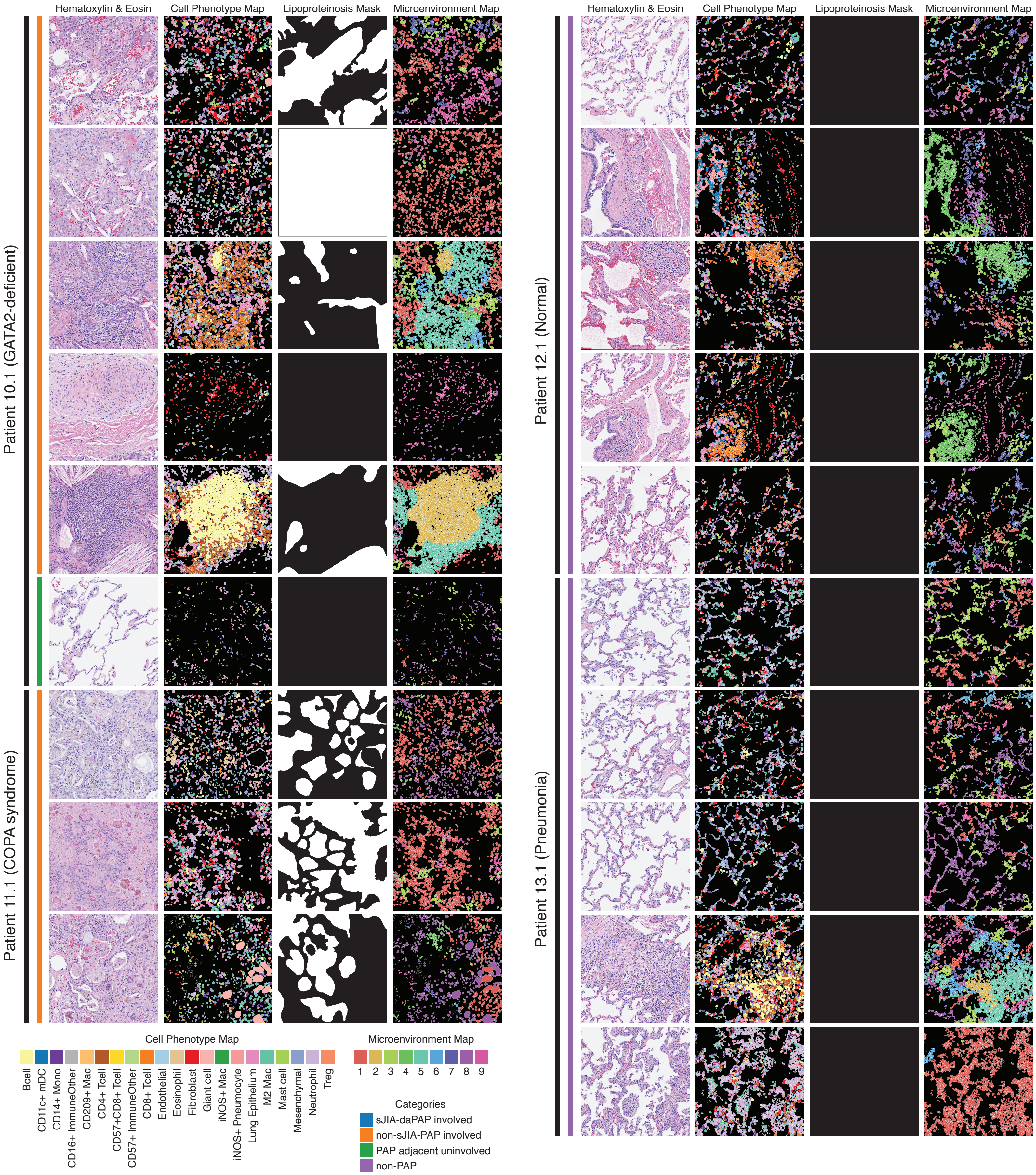

Hematoxylin & Eosin
Cell Phenotype Map
Lipoproteinosis Mask
Microenvironment Map
Patient 10.1 (GATA2-deficient)
Patient 11.1 (COPA syndrome)
Patient 12.1 (Normal)
Patient 13.1 (Pneumonia)
Cell Phenotype Map
Bcell
CD11c+ mDC
CD14+ Mono
CD16+ ImmuneOther
CD209+ Mac
CD4+ Tcell
CD57+CD8+ Tcell
CD57+ ImmuneOther
CD8+ Tcell
Endothelial
Eosinophil
Fibroblast
Giant cell
iNOS+ Mac
iNOS+ Pneumocyte
Lung Epithelium
M2 Mac
Mast cell
Mesenchymal
Neutrophil
Treg
Microenvironment Map
1 2 3 4 5 6 7 8 9
Categories
sJIA-daPAP involved
non-sJIA-PAP involved
PAP adjacent uninvolved
non-PAP

**Figure S2: Cellular and regional annotations of the study FOVs.** Histological image (first column; serial section), cell phenotype map (second column), lipoproteinosis mask (third column), and microenvironment map (fourth column) for each FOV (each row) acquired as part of this study. The FOVs are in ascending order, grouped by patient and specimen, and labeled with their respective study category.

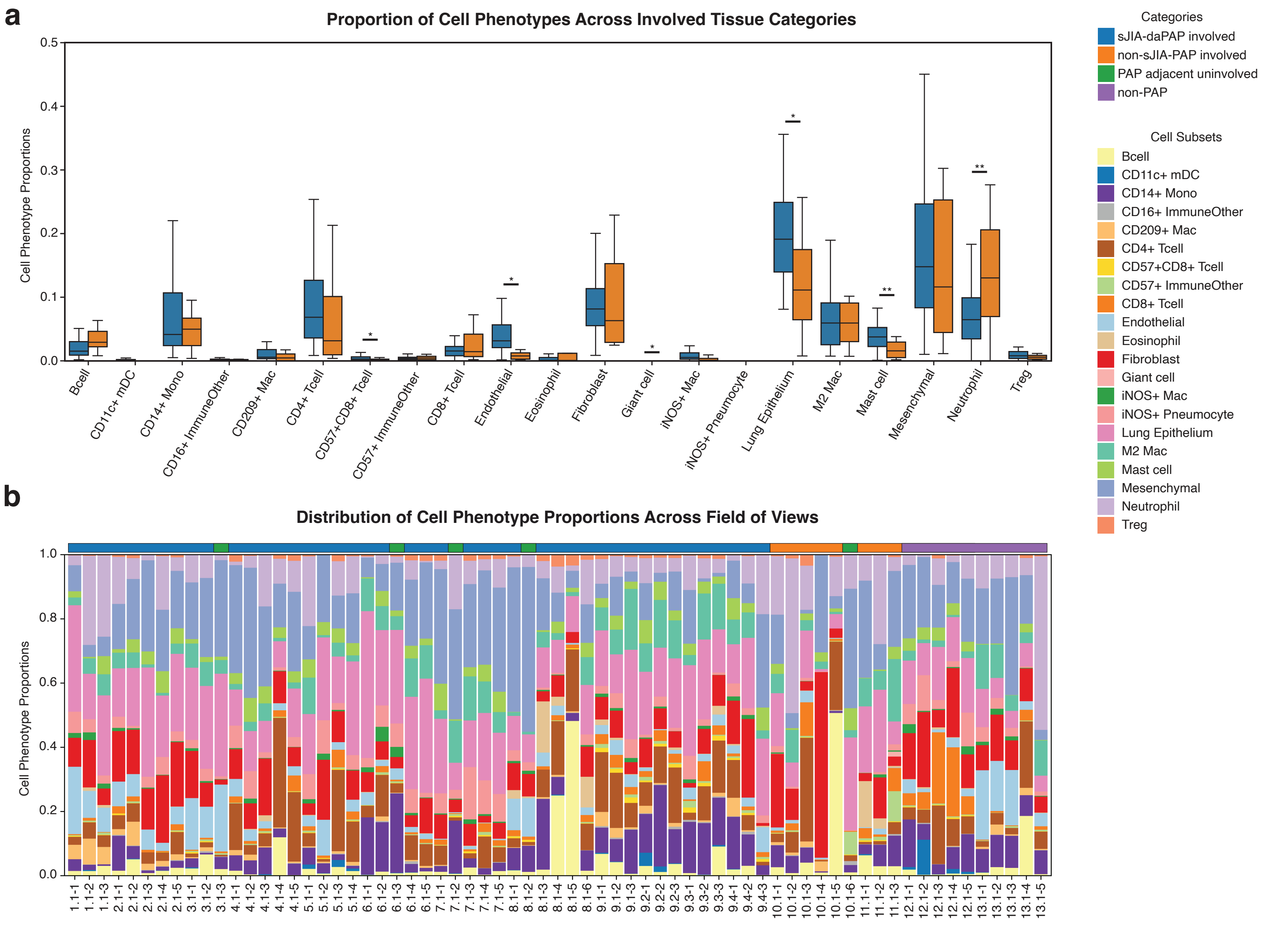
a
Proportion of Cell Phenotypes Across Involved Tissue Categories
Cell Phenotype Proportions
0.0
0.1
0.2
0.3
0.4
0.5
*
**
Bcell
CD11c+ mDC
CD14+ Mono
CD16+ ImmuneOther
CD209+ Mac
CD4+ Tcell
CD57+CD8+ Tcell
CD57+ ImmuneOther
CD8+ Tcell
Endothelial
Eosinophil
Fibroblast
Giant cell
iNOS+ Mac
iNOS+ Pneumocyte
Lung Epithelium
M2 Mac
Mast cell
Mesenchymal
Neutrophil
Treg
Categories
sJIA-daPAP involved
non-sJIA-PAP involved
PAP adjacent uninvolved
non-PAP
Cell Subsets
Bcell
CD11c+ mDC
CD14+ Mono
CD16+ ImmuneOther
CD209+ Mac
CD4+ Tcell
CD57+CD8+ Tcell
CD57+ ImmuneOther
CD8+ Tcell
Endothelial
Eosinophil
Fibroblast
Giant cell
iNOS+ Mac
iNOS+ Pneumocyte
Lung Epithelium
M2 Mac
Mast cell
Mesenchymal
Neutrophil
Treg
b
Distribution of Cell Phenotype Proportions Across Field of Views
Cell Phenotype Proportions
0.0
0.2
0.4
0.6
0.8
1.0
1.1-1 1.1-2 1.1-3 2.1-1 2.1-2 2.1-3 2.1-4 2.1-5 3.1-1 3.1-2 3.1-3 4.1-1 4.1-2 4.1-3 4.1-4 4.1-5 5.1-1 5.1-2 5.1-3 5.1-4 6.1-1 6.1-2 6.1-3 6.1-4 6.1-5 7.1-1 7.1-2 7.1-3 7.1-4 7.1-5 8.1-1 8.1-2 8.1-3 8.1-4 8.1-5 8.1-6 9.1-1 9.1-2 9.1-3 9.2-1 9.2-2 9.2-3 9.3-1 9.3-2 9.3-3 9.4-1 9.4-2 9.4-3 10.1-1 10.1-2 10.1-3 10.1-4 10.1-5 10.1-6 11.1-1 11.1-2 11.1-3 12.1-1 12.1-2 12.1-3 12.1-4 12.1-5 13.1-1 13.1-2 13.1-3 13.1-4 13.1-5

**Figure S3: Cellular composition of study samples.** Proportion of each microenvironment per acquired FOV across study categories and non-PAP samples. Cell subsets are ordered alphabetically, starting from zero.

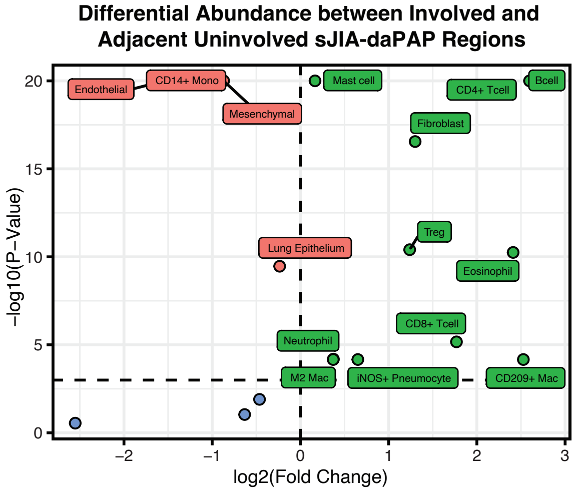

Differential Abundance between Involved and Adjacent Uninvolved sJIA-daPAP Regions
Endothelial
CD14+ Mono
Mesenchymal
Mast cell
CD4+ Tcell
Bcell
Fibroblast
Treg
Lung Epithelium
Eosinophil
CD8+ Tcell
Neutrophil
M2 Mac
iNOS+ Pneumocyte
CD209+ Mac
−log10(P−Value)
log2(Fold Change)
20
15
10
5
0
−2
−1
0
1
2
3

**Figure S4: Differential abundance of cellular subsets between sJIA-daPAP involved and adjacent uninvolved regions.** Significance threshold of absolute value of log2 fold change > 0 and adjusted p-value < 0.05, with Benjamini/Hochberg multiple hypothesis correction applied.

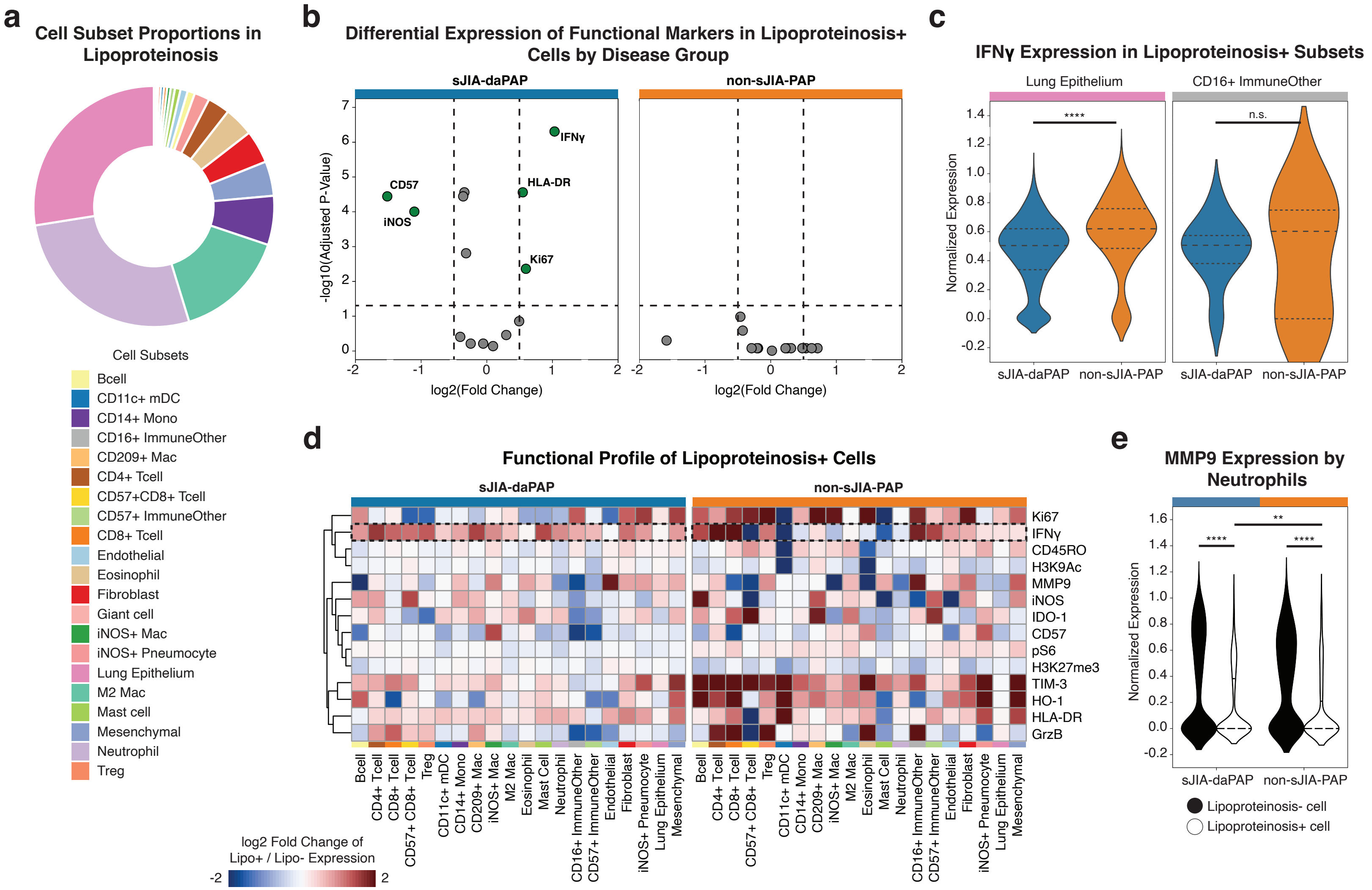

a
Cell Subset Proportions in Lipoproteinosis
Cell Subsets
Bcell
CD11c+ mDC
CD14+ Mono
CD16+ ImmuneOther
CD209+ Mac
CD4+ Tcell
CD57+CD8+ Tcell
CD57+ ImmuneOther
CD8+ Tcell
Endothelial
Eosinophil
Fibroblast
Giant cell
iNOS+ Mac
iNOS+ Pneumocyte
Lung Epithelium
M2 Mac
Mast cell
Mesenchymal
Neutrophil
Treg
b
Differential Expression of Functional Markers in Lipoproteinosis+ Cells by Disease Group
sJIA-daPAP
non-sJIA-PAP
-log10(Adjusted P-Value)
log2(Fold Change)
IFNγ
HLA-DR
CD57
iNOS
Ki67
c
IFNγ Expression in Lipoproteinosis+ Subsets
Lung Epithelium
CD16+ ImmuneOther
Normalized Expression
****
n.s.
sJIA-daPAP
non-sJIA-PAP
d
Functional Profile of Lipoproteinosis+ Cells
sJIA-daPAP
non-sJIA-PAP
Ki67
IFNγ
CD45RO
H3K9Ac
MMP9
iNOS
IDO-1
CD57
pS6
H3K27me3
TIM-3
HO-1
HLA-DR
GrzB
Bcell
CD4+ Tcell
CD8+ Tcell
CD57+ CD8+ Tcell
Treg
CD11c+ mDC
CD14+ Mono
CD209+ Mac
iNOS+ Mac
M2 Mac
Eosinophil
Mast Cell
Neutrophil
CD16+ ImmuneOther
CD57+ ImmuneOther
Endothelial
Fibroblast
iNOS+ Pneumocyte
Lung Epithelium
Mesenchymal
log2 Fold Change of Lipo+ / Lipo- Expression
-2
2
e
MMP9 Expression by Neutrophils
Normalized Expression
****
**
sJIA-daPAP
non-sJIA-PAP
Lipoproteinosis- cell
Lipoproteinosis+ cell

**Figure S5: Functional profile of cells within lipoproteinosis varies between sJIA-daPAP and non-sJIA-PAP samples. (a)** Cell subset proportions among cells localized within lipoproteinosis regions. **(b)** Differential expression of lipoproteinosis+ cells, broken down by study category. Significance threshold of absolute value of log2FC > 0.5 and adjusted p-value of 0.05, with Benjamini/Hochberg multiple hypothesis correction applied. **(c)** Comparison of IFN$\gamma$ expression by cells subsets between sJIA-daPAP and non-sJIA-PAP samples. **(d)** Heatmap of the log2 fold change of the mean normalized expression from lipoproteinosis+ cells over that from lipoproteinosis- cells for all the functional markers. Columns are ordered by phenotype group (lymphocytes, myeloid cells, granulocytes, immune "other", and non-immune), whereas rows are hierarchically clustered (Euclidean distance, average linkage). **(e)** Comparison of MMP9 expression by neutrophils between lipoproteinosis+ and lipoproteinosis- regions within sJIA-daPAP and non-sJIA-PAP samples. All P values were calculated with a Student's t-test (two tailed) (**P<0.01, ****P < 0.0001).

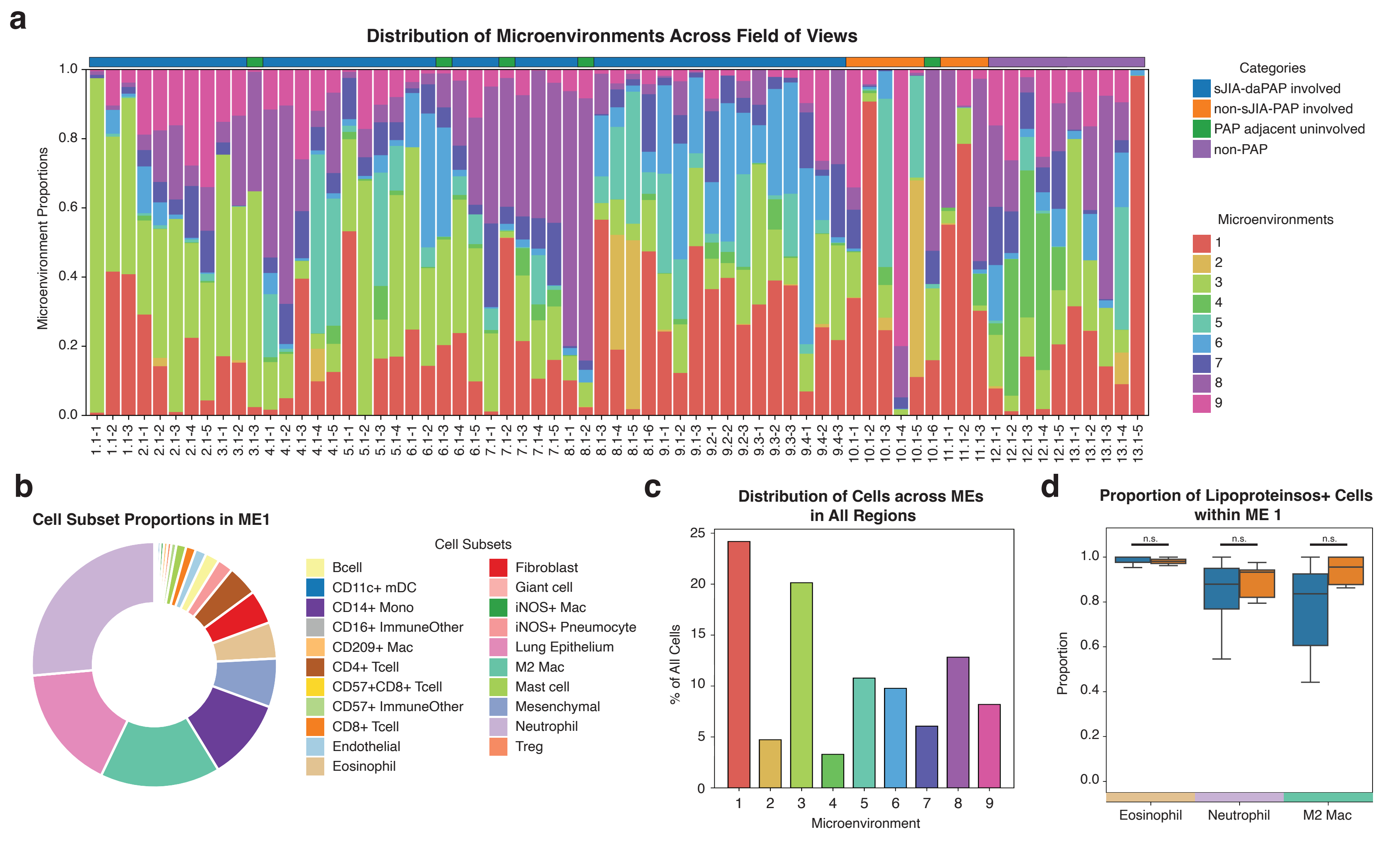
a
Distribution of Microenvironments Across Field of Views
Microenvironment Proportions
0.0
0.2
0.4
0.6
0.8
1.0
1.1-1 1.1-2 1.1-3 2.1-1 2.1-2 2.1-3 2.1-4 2.1-5 3.1-1 3.1-2 3.1-3 4.1-1 4.1-2 4.1-3 4.1-4 4.1-5 5.1-1 5.1-2 5.1-3 5.1-4 6.1-1 6.1-2 6.1-3 6.1-4 6.1-5 7.1-1 7.1-2 7.1-3 7.1-4 7.1-5 8.1-1 8.1-2 8.1-3 8.1-4 8.1-5 8.1-6 9.1-1 9.1-2 9.1-3 9.2-1 9.2-2 9.2-3 9.3-1 9.3-2 9.3-3 9.4-1 9.4-2 9.4-3 10.1-1 10.1-2 10.1-3 10.1-4 10.1-5 10.1-6 11.1-1 11.1-2 11.1-3 12.1-1 12.1-2 12.1-3 12.1-4 12.1-5 13.1-1 13.1-2 13.1-3 13.1-4 13.1-5
Categories
sJIA-daPAP involved
non-sJIA-PAP involved
PAP adjacent uninvolved
non-PAP
Microenvironments
1
2
3
4
5
6
7
8
9
b
Cell Subset Proportions in ME1
Cell Subsets
Bcell
CD11c+ mDC
CD14+ Mono
CD16+ ImmuneOther
CD209+ Mac
CD4+ Tcell
CD57+CD8+ Tcell
CD57+ ImmuneOther
CD8+ Tcell
Endothelial
Eosinophil
Fibroblast
Giant cell
iNOS+ Mac
iNOS+ Pneumocyte
Lung Epithelium
M2 Mac
Mast cell
Mesenchymal
Neutrophil
Treg
c
Distribution of Cells across MEs in All Regions
% of All Cells
0
5
10
15
20
25
1 2 3 4 5 6 7 8 9
Microenvironment
d
Proportion of Lipoproteinsos+ Cells within ME 1
Proportion
0.0
0.2
0.4
0.6
0.8
1.0
n.s.
n.s.
n.s.
Eosinophil
Neutrophil
M2 Mac

**Figure S6: Microenvironment modeling across PAP-involved samples. (a)** Proportion of each microenvironment per acquired FOV across study categories and non-PAP samples. Microenvironments are ordered numerically, starting from zero. **(b)** Cell subset proportions among cells affiliated with ME1. **(c)** Percent of cells associated with each microenvironment across all three study categories: sJIA-daPAP, non-sJIA-PAP, and adjacent uninvolved samples. **(d)** Frequency distribution of lipoproteinosis+ eosinophils, neutrophils, and M2 macrophages in ME1 out of all MEs between sJIA-daPAP and non-sJIA-PAP FOVs.

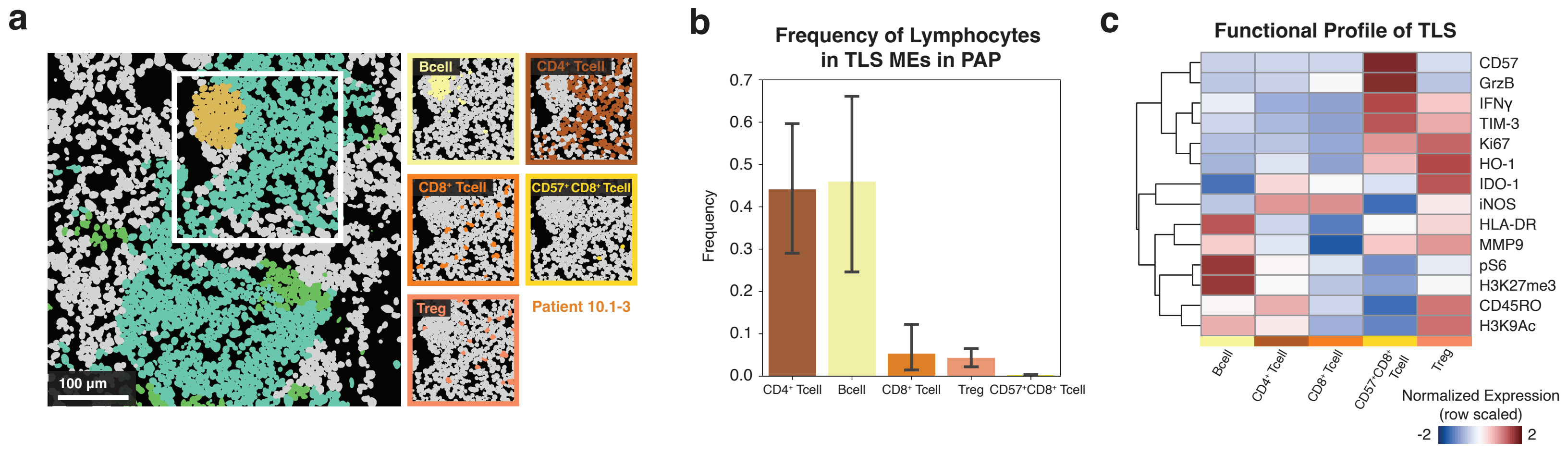
a
Bcell
CD4+ Tcell
CD8+ Tcell
CD57+CD8+ Tcell
Treg
Patient 10.1-3
100 μm
b
Frequency of Lymphocytes in TLS MEs in PAP
Frequency
0.0
0.1
0.2
0.3
0.4
0.5
0.6
0.7
CD4+ Tcell
Bcell
CD8+ Tcell
Treg
CD57+CD8+ Tcell
c
Functional Profile of TLS
CD57
GrzB
IFNγ
TIM-3
Ki67
HO-1
IDO-1
iNOS
HLA-DR
MMP9
pS6
H3K27me3
CD45RO
H3K9Ac
Bcell
CD4+ Tcell
CD8+ Tcell
CD57+CD8+ Tcell
Treg
Normalized Expression (row scaled)
-2
2

**Figure S7: Tertiary lymphoid structures are features in PAP-diseased tissues. (a)** Representative FOV with a tertiary lymphoid structure (TLS), colored by MEs associated with TLSs (left). On right, CPMs of inset from left, colored by lymphocyte subset. **(b)** Frequency of each lymphocyte subset present in MEs 2, 4, and 5 in the FOVs where a TLS was identified. **(c)** Functional profile of the lymphocyte subsets present in the TLSs. Rows are mean normalized. Columns are ordered alphabetically, whereas rows are ordered with hierarchical clustering (Euclidean distance, average linkage).
